# RNA structure through multidimensional chemical mapping

**DOI:** 10.1101/038679

**Authors:** Siqi Tian, Rhiju Das

## Abstract

The discoveries of myriad non-coding RNA molecules, each transiting through multiple flexible states in cells or virions, present major challenges for structure determination. Advances in high-throughput chemical mapping give new routes for characterizing entire transcriptomes *in vivo,* but the resulting one-dimensional data generally remain too information-poor to allow accurate *de novo* structure determination. Multidimensional chemical mapping (MCM) methods seek to address this challenge. Mutate-and-map (M^2^), RNA interaction groups by mutational profiling (RING-MaP and MaP-2D analysis) and multiplexed ·OH cleavage analysis (MOHCA) measure how the chemical reactivities of every nucleotide in an RNA molecule change in response to modifications at every other nucleotide. A growing body of *in vitro* blind tests and compensatory mutation/rescue experiments indicate that MCM methods give consistently accurate secondary structures and global tertiary structures for ribozymes, ribosomal domains and ligand-bound riboswitch aptamers up to two hundred nucleotides in length. Importantly, MCM analyses provide detailed information on structurally heterogeneous RNA states, such as ligand-free riboswitches, that are functionally important but difficult to resolve with other approaches. The sequencing requirements of currently available MCM protocols scale at least quadratically with RNA length, precluding general application to transcriptomes or viral genomes at present. We propose a modify-crosslink-map expansion to overcome this and other current limitations to resolving the *in vivo* ‘RNA structurome’.

## 1. Introduction

RNA molecules underlie many of the core processes of life. RNA’s biological roles include catalysis of peptide bond formation and deciphering the genetic code in all living systems; elaborate alternative splicing of RNA messages in different tissues during metazoan development and evolution; and packaging, replication, and processing of pervasive parasitic elements, including viruses and retrotransposons [see (Gesteland et al., 2006) and references therein]. Even as the RNAs involved in these processes have been under intense investigation, a vast number of additional RNA molecules are being discovered in genomic segments that do not code for proteins but appear to be transcribed and processed in a regulated manner; see (Amaral et al., 2008; Eddy, 2014; Qureshi & Mehler, 2012) and references therein. Understanding whether, when, and how these RNA molecules functionally impact complex organisms is a major current challenge in biology.

Well-studied ‘RNA machines’ such as the ribosome and the spliceosome form and interconvert between intricate three-dimensional structures as they sense and respond to their protein, nucleic acid, and small molecule partners. It is possible that some or many of the newly discovered non-coding RNA molecules may transit through such functional structures and even interact to form an extended RNA machine (Amaral et al., 2008). However, it is also possible that most non-coding RNAs harbor sparse or no regions that form functional structures. In either case, these possibilities are, for the most part, untested. On one hand, structure determination methods that achieve high resolution are growing in power and applicability, with recent improvements in cryo-electron microscopy achieving near-atomic-resolution models for RNA complexes extracted from living cells (Amunts et al., 2014; Greber et al., 2014; Hang et al., 2015; Nguyen et al., 2015). On the other hand, these methods, along with crystallography and NMR approaches, continue to face challenges in RNAs that form non-compact states, form multiple structures, bind a heterogeneous complement of partners, or that have large unstructured regions.

In contrast to high-resolution methods, chemical mapping (also called ‘footprinting’, ‘chemical probing’, or ‘structure mapping’) experiments can be applied to most RNAs under most solution conditions, including molecules that form heterogeneous, flexible structures or molecules functioning in their native cellular or viral milieu. Chemical mapping methods mark nucleotides that are accessible to chemical attack. Such reactivity is typically correlated to nucleotide solvent accessibility or flexibility, key features of RNA structure. As these techniques are read out by nucleic acid sequencing, chemical mapping methods have undergone accelerations over the last decade as sequencing technologies have rapidly advanced, enabling characterization of RNA chemical accessibilities of entire transcriptomes *in vivo;* see, e.g., and refs. therein. These experiments raise the prospect of nucleotide-resolution structural portraits of all RNAs being transcribed in an organism – the ‘RNA structurome’. Nevertheless, when tested through independent experiments, *de novo* models derived from chemical mapping and computational modeling have not always given consistently accurate structures, even on small domains folded into well-defined states and probed *in vitro* (Deigan et al., 2009; Kladwang et al., 2011c; Leonard et al., 2013; Tian et al., 2014). These issues can be traced to the poor information content of chemical mapping measurements, which typically give single or few measurements per nucleotide, compared to high-resolution technologies such as crystallography, NMR, or cryo-electron microscopy, which can return data sets with ten or more measurements per nucleotide.

Multidimensional chemical mapping (MCM) techniques were proposed recently to address the limited information content of conventional chemical mapping data (Das et al., 2008; Kladwang & Das, 2010). MCM methods seek to determine not just chemical reactivities at each nucleotide but also how these reactivities are affected by systematic perturbations – nucleotide mutations, chemical modifications, or radical source attachments – at every other nucleotide (Figure 1). Analogous to multidimensional forms of NMR spectroscopy, such multidimensional chemical data were hypothesized to give sufficient constraints to accurately model RNA secondary structure and tertiary structure at nucleotide resolution and to give detailed empirical information on heterogenous ensembles. If successful, MCM would provide a needed ‘front-line’ technique for inferring RNA structure: structured domains of long RNA transcripts could be rapidly defined and visualized from *in vivo* experiments. If a domain interconverts between multiple structural states, such states could be further parsed and separately stabilized through mutation, again with rapid nucleotide-resolution tests by MCM. After initial MCM-guided analysis, these domains would then become candidates for more detailed biochemical analysis, including discovery of protein partners; functional analysis through *in vivo* mutation and epistasis experiments; and detailed structural dissection through high-resolution techniques, such as crystallography and cryo-electron microscopy. However, prior to investing efforts into developing an MCM-initialized pipeline, it has been necessary to test the hypothesis that MCM methods will actually produce sufficient information to model RNA structures *de novo.* The purpose of this article is to review recent studies on model systems and newly discovered RNAs that have evaluated this basic hypothesis, setting the stage for *in vivo* expansions.

**Figure 1.**
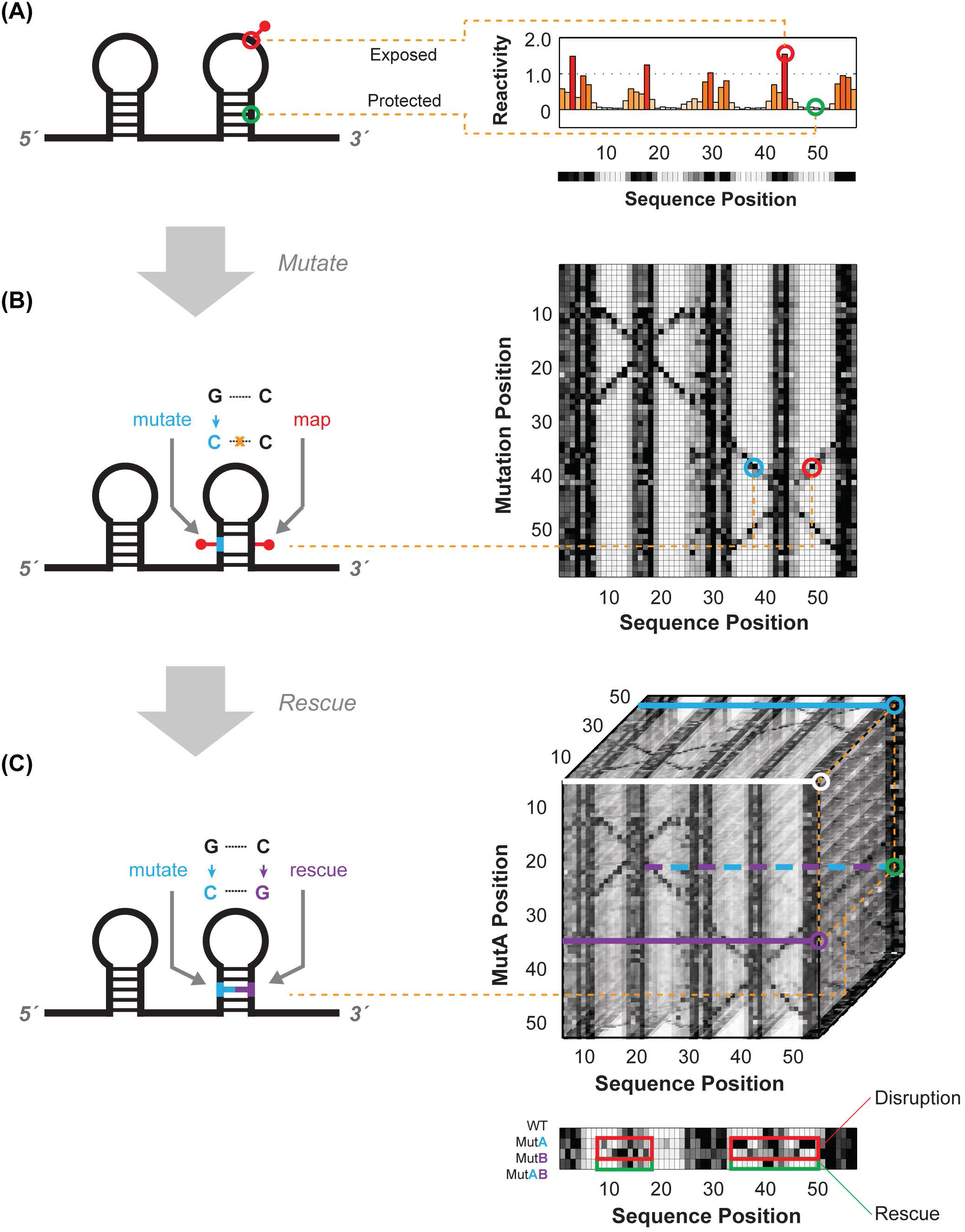
Schematics for multidimensional expansions of chemical mapping to infer RNA structure. (A). Schematic of one-dimensional (1D) chemical mapping and simulated reactivity profile. The red pin illustrates a chemical modification event on an exposed (non base-pairing) nucleotide. The red and green circles highlight a reactive (exposed) and unreactive (protected) nucleotide, respectively. (B). Schematic of two-dimensional (2D) chemical mapping through the mutate-and-map (M^2^) strategy. A sequence mutation (cyan) breaks a base-pair, exposing both itself and its partner (red), resulting in measurable increases in chemical reactivity at the partner (right). On a full dataset with mutations made separately at every position (right), a diagonal feature should trace perturbations near each single mutation position, while cross-diagonal features should report individual residues released upon mutation of their pairing partners. (C). Schematic of 3D chemical mapping. When all double mutants are chemically mapped, the entire dataset would fill a cube (mutate-mutate-map, M^3^, right). In practice, a smaller set of single and compensatory double mutations can target particular base-pair hypotheses. A quartet of chemical mapping profiles (WT, MutA, MutB, and MutAB) illustrates mutate-map-rescue (M^2^R, bottom). Here, perturbations that occur upon single mutations (at base pair partners, in MutA; or delocalized changes, in MutB; outlined in red) are rescued upon concomitant double mutation (outlined in green, MutAB). In all panels, simulated data are shown to illustrate concepts; see subsequent figures for experimental data. Orange dotted lines connect specific nucleotides or nucleotide pairs in RNA (left) to corresponding positions in multidimensional data (right).

The organization of the review is as follows. The next section (Section 2) briefly summarizes recent improvements to conventional 1D chemical mapping methods and their current limitations, motivating the development of MCM. Section 3 describes the best-tested MCM approach, the mutate-and-map (M^2^) technique, including its conception, its experimental evaluation, and a recent acceleration through mutational profiling (MaP). Section 4 describes and evaluates a second MCM method hypothesized to complement M^2^ with longer-distance data needed for three-dimensional modeling, called multiplexed ·OH cleavage analysis (MOHCA). Section 5 illustrates first applications of MCM to characterize RNA states with significant secondary structure or tertiary structure heterogeneity, including ligand-free riboswitch states intractable to other high-throughput methods. Section 6 summarizes current challenges in bringing MCM to bear on RNA transcripts longer than a few hundred nucleotides, especially within their biological milieu. These challenges include not only technical issues in making comprehensive nucleotide-level perturbations to cellular RNAs but also a more fundamental problem in how MCM sequencing costs scale with RNA length. A modify-crosslink-map protocol – not yet put into practice – is proposed to solve these problems. A summary of the MCM methods reviewed herein is presented in Table 1. Conclusions in the review make use of publicly available data deposited in the RNA Mapping Database (Cordero et al., 2012b); accession IDs are listed in figure legends. Section 7 summarizes the review.

**Table 1.** Multidimensional chemical mapping methods for RNA structure characterization.

| Method | References | Perturbation |  | Data acquired for perturbation |  | Total no. data<sup>a</sup> |
| --- | --- | --- | --- | --- | --- | --- |
|  |  | Type | Number<sup>a</sup> | Type | Number<sup>a</sup> |  |
| Mutate-and-map ($M^2$) | (Kladwang et al., 2011a; Kladwang & Das, 2010; Kladwang et al., 2011b) | Mutation, encoded in DNA template | $O(N)$ | Modification/cleavage sites | $O(N)$ | $O(N^2)$ |
| Multiplexed $\bullet$OH cleavage analysis (MOHCA) | (Cheng et al., 2015b; Das et al., 2008) | Fe(II) chelate introduced during transcription | $O(N)$ | RNA cleavage sites | $O(N)$ | $O(N^2)$ |
| RNA Interaction Groups by Mutational Profiling (RING-MaP) and MaP2D | (Homan et al., 2014) | Covalent modification by solution probe | $O(N)$ | Modification sites at other nucleotides | $O(N)$ | $O(N^2)$ |
| Mutate-map-rescue ($M^2R$) | (Tian et al., 2014) | Single/double mutations, encoded in DNA template | $O(N)$ | Modification/cleavage sites | $O(N)$ | $O(N^2)$ |
| Mutate-mutate-map ($M^3$) | Unpublished | All single & double mutations | $O(N^2)$ | Modification/cleavage sites | $O(N)$ | $O(N^3)$ |
| Modify-crosslink-map (MXM) | Proposed herein | Covalent modification by solution probe | $O(N)$ | Modification sites in cross-linked fragments | $O(\log N)$ | $O(N \log N)$ |

<sup>a</sup> $N$ is the number of nucleotides in the RNA.

## 2. Prelude: one-dimensional RNA chemical mapping

RNA structure has been empirically probed by ‘one-dimensional’ chemical mapping experiments for more than three decades. As a classic example, dimethyl sulfate (DMS) was tested as a structural probe almost immediately after its development for nucleic acid sequencing (Peattie & Gilbert, 1980). DMS remains in use to methylate the N1/N3 atoms of A/C nucleobases that have their Watson-Crick edges exposed to solution. Modification by DMS thus reports that a nucleotide is not engaged in a Watson-Crick pair in the secondary structure (Cordero et al., 2012a; Tijerina et al., 2007). Chemical modification by DMS or other probes can be rapidly read out at every nucleotide of an RNA through primer extension reactions that terminate immediately 3’ to the modified bases, followed by electrophoresis or next-generation sequencing of the resulting cDNA products. The currently available set of chemical and enzymatic probes of RNA structure and several methodological accelerations have been described in several recent reviews (Eddy, 2014; Kwok et al., 2015; Weeks, 2010) and these methods continue to be advanced; see, e.g., (Kielpinski & Vinther, 2014b; Poulsen et al., 2015; Spitale et al., 2015).

Chemical mapping measurements provide <u>one</u>-<u>d</u>imensional (1D) profiles of structure along entire transcripts (Figure 1A). These data, even in their raw form, can yield biological insights. For example, in recent transcriptome-wide studies, comparisons of *in vitro* and *in vivo* averaged structural accessibilities over numerous transcripts have illuminated the pervasive remodeling of RNA structure in cells, presumably by protein partners. Nevertheless, *de novo* structure determination from chemical mapping data has been more challenging. The protection of a given nucleotide from chemical modification does not directly reveal the nucleotide’s interaction partner, which may be any of the other protected nucleotides in the transcript or, in the case of multi-molecular complexes, other molecular partners. Chemical cross-linking approaches can pinpoint pairing partners but give sparse data (few cross-links per molecule) and, not infrequently, artifacts that have strongly distorted 3D structure models; see, e.g., studies on tRNA, ribosomes, group II introns, and the spliceosome (Anokhina et al., 2013; Dai et al., 2008; Hang et al., 2015; Levitt, 1969; Robart et al., 2014; Sergiev et al., 2001; Whirl-Carrillo et al., 2002). The information content of chemical mapping is therefore low. Until recently, expert intuition and *ad hoc* manual comparison of chemical mapping data with phylogenetic information and computational methods have been necessary to integrate chemical data into structure models, sometimes leading to significant errors (Anokhina et al., 2013; Dai et al., 2008; Deigan et al., 2009; Hang et al., 2015; Levitt, 1969; Robart et al., 2014; Sergiev et al., 2001; Tian et al., 2014; Whirl-Carrillo et al., 2002).

Several studies suggested that direct integration of one-dimensional chemical mapping data into energy-optimizing computational algorithms as ‘pseudoenergies’ would enable automated *de novo* secondary structure determination with high accuracy. There have been promising results on several model RNAs of known structure, including large molecules such as the 1542-nucleotide *E. coli* 16S ribosomal RNA (Deigan et al., 2009; Hajdin et al., 2013; Rice et al., 2014). However, the general level of accuracy of these techniques for new RNAs has been questioned (Kladwang et al., 2011c; Sukosd et al., 2013; Tian et al., 2014). For example, reanalysis of a model based on selective 2’-OH acylation by primer extension (SHAPE) of the 9173-nucleotide HIV-1 RNA genome (Watts et al., 2009) suggested that more than half of the presented helices were not well-determined (Kladwang et al., 2011c), and subsequent work, including both experimental and computational improvements, have significantly revised these uncertain regions (Pollom et al., 2013; Siegfried et al., 2014; Sukosd et al., 2015). The debate over whether these methods produce acceptable structure accuracies continues (Deigan et al., 2009; Eddy, 2014; Kladwang et al., 2011c; Leonard et al., 2013; Rice et al., 2014; Sukosd et al., 2013; Tian et al., 2014) and will not be reviewed in detail here. There is general agreement, however, on some points. First, combination of chemical mapping data with automated algorithms provides more predictive power and more reproducible results than using either method separately. Second, these methods face limitations when applied to RNAs that form significant tertiary structure, that form complexes with proteins or other molecular partners, or that populate multiple states (Leonard et al., 2013). These issues preclude the application of one-dimensional chemical mapping to automated RNA domain structure detection – much less *de novo* structure determination – in many biological contexts of interest.

## 3. Mutate-and-map for 2D structure

### 3.1 Mutate-and-map (M^2^) concept

The secondary structure and tertiary interactions of an RNA structure are defined by a list of which nucleotides come together to form Watson-Crick base pairs or non-canonical interactions. As noted above, conventional one-dimensional chemical mapping constrains but does not directly return this list of pairings. In particular, the data do not directly report the pairing partner(s) of each protected nucleotide (Figure 1A).

The <u>M</u>utate-and-<u>M</u>ap (M^2^) approach was proposed in 2010 as a potentially general experimental route to resolve the ambiguity of RNA pairing partners (Kladwang & Das, 2010). The proposal was conceptually straightforward: If two nucleotides are paired in the RNA structure, mutation of one nucleotide might ‘release’ both partners, producing localized changes observable in single-nucleotide-resolution chemical mapping profiles. The proposed effect is illustrated in Figure 1B, and was supported by observations in prior mutational studies on group I introns (Garcia & Weeks, 2004; Pyle et al., 1992). In general, disruption by a single mutation might not give precise release of partners but instead produce global unfolding of the RNA, localized unfolding of stems, or refolding of the RNA into an alternative structure. Fortunately, chemical mapping data would still discriminate between these scenarios based on the number and pattern of nucleotides with perturbed chemical reactivity. If even a subset of mutations give the desired pinpointed disruption of partners, this would provide strong information on RNA structure. However, at the time of the proposal, it was unclear if such an informative subset of mutations would generally be found in structured RNAs.

### 3.2 Proof-of-concept in designed systems

The M^2^ proposal motivated the development of methods to synthesize variants mutating every position in a nucleic acid sequence, analogous to alanine scanning in proteins but not carried out routinely in RNA biochemical studies. The proposal also motivated advances in high-throughput protocols for chemical mapping of these variants, replacing radioactive labeling of primers and slab gel electrophoresis with fluorescent readouts and capillary electrophoresis instruments developed for Sanger sequencing (Kladwang et al., 2011a; Mitra et al., 2008; Yoon et al., 2011). These accelerations now allow M^2^ measurements to be carried out and analyzed in two days, after receipt of automatically designed primers for template assembly from commercial DNA companies (Cordero et al., 2014; Lee et al., 2015; Tian et al., 2015).

Proof-of-concept experiments for M^2^ were encouraging. A first study was carried out on a 20 base-pair DNA/RNA hybrid helix (Kladwang & Das, 2010). This X-20/H-20 system was chosen since every possible single-nucleotide mutation and deletion to the DNA could be ordered without further processing, and the RNA’s DMS modification profile could be mapped with gel and capillary electrophoresis readouts. Visualization of the raw data showed ‘punctate’ events marking 15 of the 17 base pairs involving an A or C (the nucleotides visible to DMS read out by primer extension) on the RNA strand (outlined in orange, cyan, and green outlines; Figure 2A). Inferring these base pairs did not require visual inspection but could also be captured by an automated algorithm. The algorithm was based on Z-scores, the number of standard deviations by which reactivity at a nucleotide exceeded its mean reactivity over all constructs when a putative partner was mutated.

**Figure 2.**
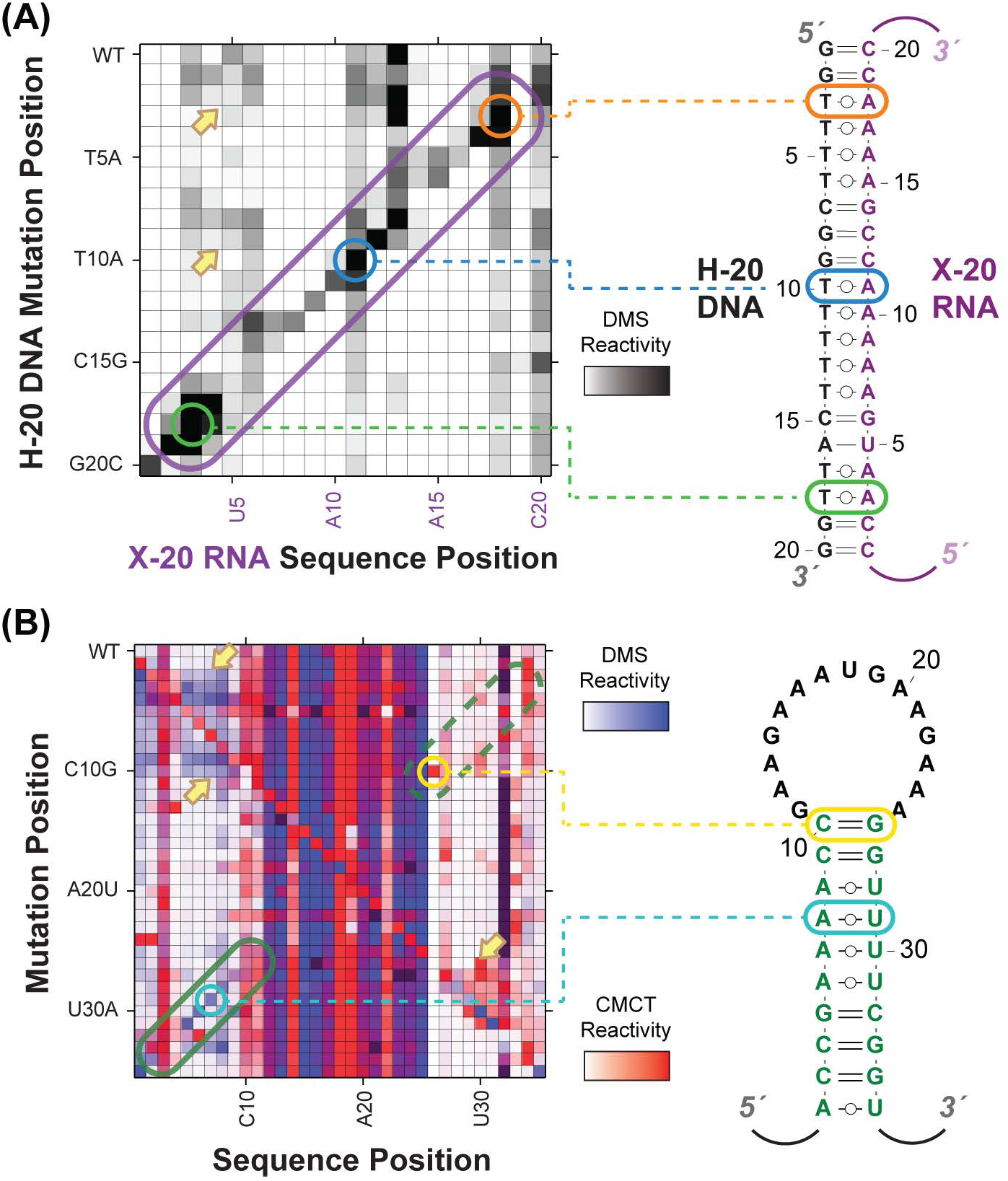
Experimental proof-of-concepts of the M^2^ methodology. (A). Experimental M^2^ measurements (left) and secondary structure (right) of a H-20/X-20 DNA/RNA hybrid construct (Kladwang & Das, 2010). Single mutations of the H-20 DNA result in mismatches in the hybrid helix, exposing nucleotides in the X-20 RNA (purple) to DMS chemical modification. Purple line outlines region with expected base pair features; orange, blue, and green circles highlight a few strong features that correspond to expected base pairs. (B). M^2^ data and secondary structure of a MedLoop test RNA (Kladwang et al., 2011a). The test helix is designed to be mostly A/C on one side and U/G on the other. DMS (blue) and CMCT (red) M^2^ datasets are overlaid. Regions corresponding to expected base pairs from the step are outlined in green on the data. Yellow and cyan circles mark a few single-nucleotide features in the M^2^ data (left) that demarcate specific base pairs (right). In both (A) and (B), yellow arrows mark perturbations from mutation that extend beyond ‘punctate’ release of a single base pair and involve disruption of an entire helix. RMDB Accession IDs for datasets shown: (A). X20H20_DMS_0001; (B). MDLOOP_DMS_0002 and MDLOOP_CMC_0002.

Further experiments on a 35-nucleotide ‘Medloop’ RNA hairpin confirmed that M^2^ could be applied to infer RNA-RNA base pairs, using data from DMS, SHAPE, and CMCT, a reagent specific to exposed G and U Watson-Crick edges. In Figure 2B, perturbations near the site of each mutation and at partners are highlighted (cyan and yellow outlines). Not every mutation gave punctate release of partners. Some showed no perturbations, presumably due to replacement of the original Watson-Crick pair with a non-Watson-Crick pair; and others gave more delocalized perturbations (yellow arrows, Figure 2A-B; see Section 5 for further discussion). Some nucleotides appeared to be ‘hotspots’, becoming exposed by many different mutations (see, e.g., G27 in Figure 2B). Nevertheless, nine of the hairpin’s ten base pairs could be inferred from a sequence-independent analysis searching for punctate features. The analysis was again based on finding M^2^ features with high Z-scores; enforcing that multiple such features clustered together was important in eliminating any of the 1460 possible false positives. This study also revealed that the strongest effects were seen when mutating each nucleotide to its complement. These most informative substitutions became the default mutation set in later studies. These early results also highlighted the importance of collecting data on mutants at all sequence positions, not only to capture base pairs throughout the RNA but also to establish whether observed perturbations were significant compared to the variability of chemical reactivity at a given site, as captured in the Z-score. Overall, these data suggested that the majority of single base pairs in a non-coding RNA might be discovered through systematic and unbiased M^2^ experiments.

### 3.3 Tests on natural RNAs

After the proofs of concept above, M^2^ studies were carried out on several RNA domains drawn from biological sources. These RNAs included a benchmark of several riboswitches and ribozyme domains that had challenged prior chemical mapping approaches (Kladwang et al., 2011b), a ribosomal domain for which (one-dimensional) SHAPE-directed modeling gave a misleading structure (Tian et al., 2014), newly discovered RNA regulons in vertebrate homeobox mRNA 5’ untranslated regions (Xue et al., 2015), and molecules presented to the RNA modeling community as ‘RNA-Puzzle’ blind challenges before publication of their crystal structures (Miao et al., 2015).

#### Initial benchmark on six natural RNAs

Visual inspection of M^2^ data for an initial benchmark of six natural non-coding RNAs provided informative lessons after the previous small, artifical proof-of-concept systems (Figure 2). As hypothesized, punctate mutation-release signals appeared in the raw M^2^ data for the natural non-coding RNAs, signaling Watson-Crick base pairs occurring in these non-coding RNAs. For example, for a double-glycine riboswitch aptamer, six helices that had been predicted by expert phylogenetic analysis – but not yet confirmed by crystallography – were visible as six cross-diagonal features in raw M^2^-SHAPE data (outlined in six different colors, Figure 3A). Nevertheless, these M^2^ data sets on biological non-coding RNA domains showed fewer punctate mutation-release signals compared to the original proof-of-concept systems (Kladwang et al., 2011a; Kladwang & Das, 2010). Indeed, for some helices, all mutations tested either gave no detectable change in chemical reactivity or produced delocalized changes in chemical mapping profiles relative to the starting sequence, suggesting unfolding or refolding of entire helices (yellow, Figure 3A). Signatures for non-canonical base pairs, including those mediating tertiary contacts, were similarly delocalized (red arrows, Figure 3A); tertiary structure will be discussed in more detail in Section 4 below. This initial visual inspection indicated that the Z-score-based inference developed with artificial systems would, on its own, not allow complete secondary structure inference, much less tertiary structure inference, of natural non-coding RNAs.

**Figure 3.**
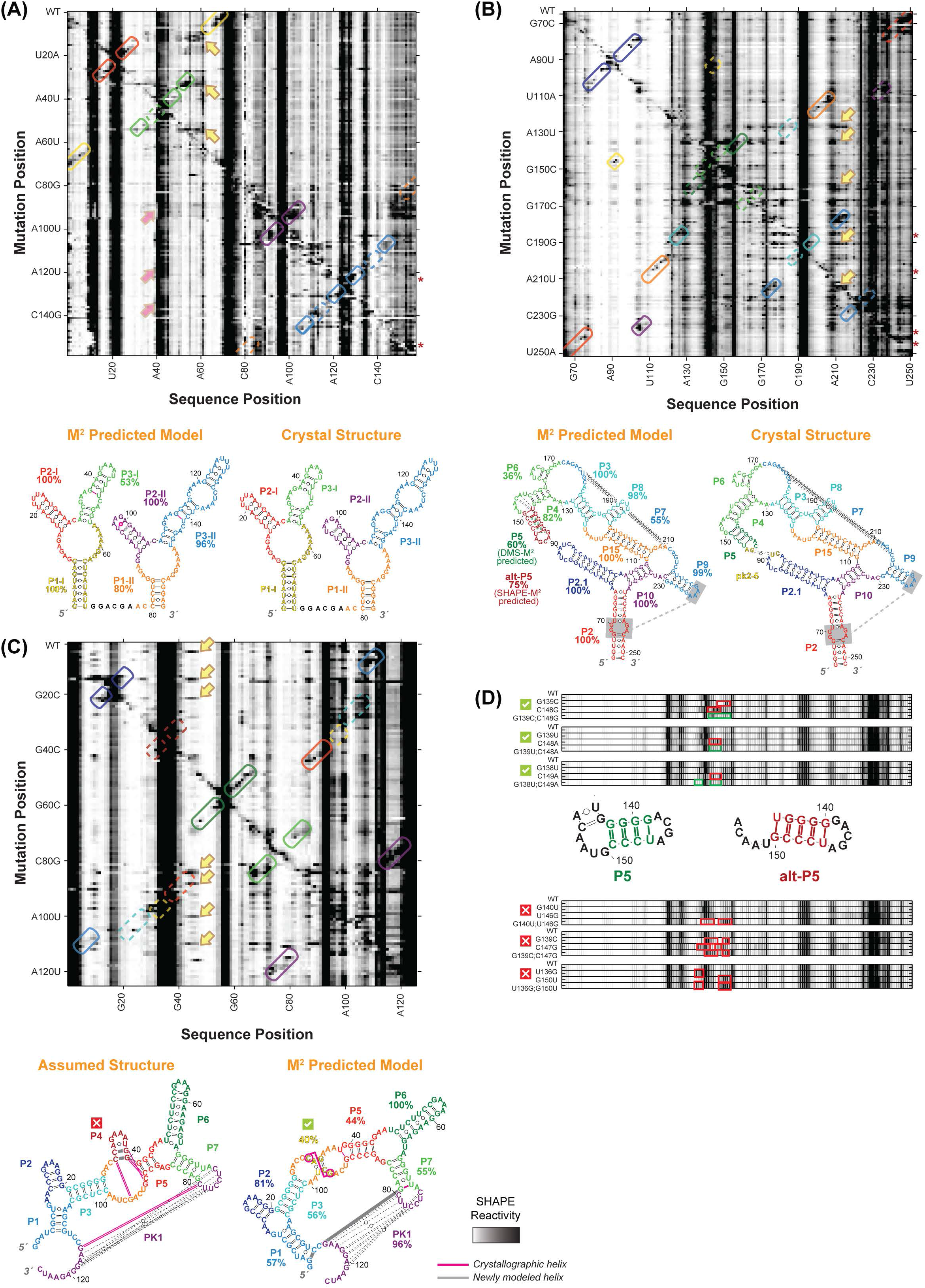
M^2^ reveals secondary structure of natural non-coding RNA domains. (A). M^2^ data and secondary structures of a double glycine riboswitch from *F. nucleatum* (Butler et al., 2011; Lipfert et al., 2007; Lipfert et al., 2010). RNA was probed in presence of 10 mM glycine. M^2^-SHAPE data are shown with helices outlined according to their assigned color. Solid outlines mark helices in which at least one mutation increases SHAPE reactivity around its expected partner and other mutations generally keep the partner protected, providing evidence for the helix; dashed outlines mark other helix locations. Red arrows mark exposure of P3-I loop upon disruption of tertiary structure that results not only from mutation of its tertiary contact partner (PI-II) but also from mutations in other helices. In secondary structures, bootstrapping confidence scores are marked under helix labels. The M^2^ predicted model using the automated Z-score analysis captured all 6 helices with > 80% bootstrapping support except for P3-I, which also has an extra base pair. (B). M^2^ data and secondary structures of the GIR1 lariat-capping ribozyme from *D. iridis,* RNA-Puzzle 5 (Miao et al., 2015). The data captured all helices and the pk2-5 tertiary contact observed in the subsequently release crystal structure (Meyer et al., 2014). Both a P5 helix (dark green) and an alternative alt-P5 (dark red), differing by a single-nucleotide register shift, were modeled by M^2^ with similar bootstrap supports. Visual inspection of M^2^-DMS [not shown; see (Miao et al., 2015)] suggested a tertiary contact involving non-canonical pairs between P9 and P2 (gray) that was indeed observed in the subsequently released crystal structure. (C). M^2^ data and secondary structures of the ydaO cyclic-di-adenosine riboswitch, RNA-Puzzle 12 (Gao & Serganov, 2014; Ren & Patel, 2014). RNA was probed in presence of 10 μM c-di-AMP. The differences of each model compared to subsequently released crystallographic structure are marked by magenta and gray lines. The secondary structure based on expert sequence analysis (left), assumed by all RNA-Puzzle modelers, included an incorrect P4 (dark red), while the M^2^ predicted model (right) correctly rearranged this region. (D). M^2^-rescue data and secondary structures of the GIR1 lariat-capping ribozyme from *D. iridis.* The discrepancy in M^2^-predicted model was resolved by M^2^-rescue data testing base-pairs in P5 and alt-P5, showing that compensatory double mutations predicted to rescue P5 succeeded in restoring the sequence’s chemical mapping profile (outlined in green) after their disruption by single mutations (outlined in red), while double mutants based on alt-P5 failed to rescue the profile. In panels (A)-(C), yellow arrows mark perturbations from mutation that involve disruption of helices or formation of alternative secondary structure. In panels (A) and (B), rows with red asterisks are mutants for which data were not acquired; to aid visual inspection, these rows have been filled in with wild type data. RMDB Accession IDs for datasets shown: (A). GLYCFN_SHP_0004; (B). RNAPZ5_1M7_0002; (C). RNAPZ12_1M7_0003; (D). unpublished result.

#### Integration with automated secondary structure prediction

The benchmark results described above (Kladwang et al., 2011b) motivated the integration of M^2^ data with well-developed secondary structure prediction methods, inspired by prior work involving one-dimensional chemical mapping (Deigan, 2009). RNAstructure and other methods predict the lowest energy (highest probability) secondary structure for an RNA sequence, given an energetic model. To guide these calculations to higher accuracy secondary structures, nucleotide pairs that gave high Z-scores in M^2^ data were assigned a proportionally strong energy bonus in RNAstructure. Across the benchmark, the resulting automatically generated secondary structures were consistently accurate, with only 1 of 185 base pairs missed, and any mispredicted base pairs occurring at only at the edges of helices (Figures 3A-3C) (Kladwang et al., 2011b). Furthermore, building on prior efforts to estimate reliability of 1D-mapping-guided secondary structures (Kladwang et al., 2011c), an analysis was developed to estimate the helix-by-helix uncertainty in M^2^-guided secondary structures, based on the recovery of each helix in ‘mock’ analyses in which the M^2^ data were randomly resampled with replacement [nonparametric boostrapping (Efron & Tibshirani, 1998)]. These analyses exposed misleading inferences from conventional chemical mapping methods (Deigan et al., 2009; Tian et al., 2014), and uncertainties in register shifts (Figure 3D, P5 vs. alt-P5) or in helices (typically short 2-bp stems) that could be further tested (see below, Section 3.4).

#### RNA-Puzzle tests

As in other areas of macromolecule modeling (Das & Baker, 2008; Fleishman et al., 2010), the strongest tests of structure prediction have been blind tests. For most of the recent blind RNA-Puzzle targets, M^2^ data were acquired and shared with all modelers during the prediction period, before crystal structures were released after modeling. These targets include two problems (the *D. iridis* lariat-capping GIR1 ribozyme and the S. *thermophilum* adenosylcobalamin riboswitch) recently summarized in the RNA-Puzzles Round II paper (Meyer et al., 2014; Miao et al., 2015; Peselis & Serganov, 2012) and four others for which crystal structures have since been reported (Ren & Patel, 2014; Suslov, 2012; Suslov et al., 2015; Trausch et al., 2015; Trausch et al., 2014).

The M^2^-based analysis has consistently achieved accurate secondary structures, including stems that are scrambled with standard computational modeling and 1D chemical mapping analysis [see, e.g., Supporting Information in (Miao et al., 2015)] and features that could not be captured by prior phylogenetic analysis (Figures 3B and 3D). For example, the precise mutation-release signals in M^2^ data revealed novel interactions for the lariat-capping GIR1 ribozyme (RNA-Puzzle 5). Mutations in nucleotide G144 and A145 exposed nucleotides C92 and U91, respectively, making apparent a P2.1/P4 pseudoknot (yellow box, Figure 3B) missed by conventional chemical mapping and by prior sequence comparisons and expert inspection (Beckert et al., 2008). The entire M^2^-derived secondary structure was accurate compared to the subsequently released crystal structure, up to edge base pairs (Figure 3B). In addition, a tertiary contact involving an A-minor interaction was detected by visual inspection of the M^2^ data; mutation of P2 sequences changed the reactivity of the apical loop of P9. These inferences enabled 3D modeling of the GIR1 ribozyme at better than 1 nm resolution (Miao et al., 2015); see also Section 4.3 below.

Further surprising results arose during automated M^2^ secondary structure modeling of RNA-Puzzle 12, the cyclic-di-adenosine monophosphate *ydaO* riboswitch from *T. tengcongensis.* Here, automated M^2^ secondary structure modeling returned a model with nearly all the stems expected from prior expert analysis of sequence conservation and covariation, including a long-range pseudoknot PK1 (Figure 3C). However, this analysis did not recover one hairpin stem P4, even though the target sequence included a GAAA tetraloop introduced to stabilize this stem (Figure 3C). During the prediction period, our group assumed this to be a failure of the M^2^ approach, and all models from our group and all other groups included P4. Nevertheless, when the crystal structure was released, the M^2^ analysis turned out to be accurate: the crystallized RNA did not show electron density for the P4 tetraloop, and the conserved nucleotides in this region formed a non-canonical internal two-way junction instead of a hairpin stem (Gao & Serganov, 2014; Ren & Patel, 2014).

Overall, the studies carried out to date on well-structured RNAs have strongly supported the M^2^ strategy. Systematic mutagenesis can be coupled to chemical mapping to yield rich structural information hidden in or missed by conventional chemical mapping data. The data by themselves allow direct single-nucleotide-resolution inference of some Watson-Crick base pairs through punctate mutation-release signals. More generally, modeling that integrates M^2^ data with state-of-the-art secondary structure prediction methods give full models of all Watson-Crick pairs. This automated M^2^ approach has been consistently accurate at nucleotide resolution for RNAs that have been challenging for prediction methods based on computational modeling, conventional one-dimensional mapping data, phylogenetic analysis, expert analysis, or combinations thereof (Kladwang et al., 2011b; Miao et al., 2015; Tian et al., 2014). These conclusions have been borne out in 12 noncoding RNAs whose structures have been solved through crystallography, including 6 RNA-Puzzle blind modeling targets.

### 3.4 Stringent tests through mutation/rescue

The majority of RNA transcripts in biological systems will not necessarily form a single well-defined structure. Thus, the tests of the M^2^ concept above, which relied on crystallization of an RNA to give ‘gold standard’ reference structures, were incomplete. The need for more general validation or falsification motivated a further expansion of the M^2^ concept to enable not only the discovery but also the incisive testing of RNA base pairs (Figure 1C).

The <u>M</u>utate-<u>M</u>ap-<u>R</u>escue (M^2^R) proposal is a high-throughput variant of compensatory rescue experiments, which have provided strong tests of Watson-Crick base pairing in nearly every well-studied RNA system, including striking examples *in vivo* (Graveley, 2005; Lehnert et al., 1996; Madhani & Guthrie, 1994; Reenan, 2005; Singh et al., 2007). In these experiments, two partners in a putative base pair are separately mutated to their complement. If concomitant introduction of these separately disruptive mutations restores the RNA’s function, the pairing is strongly supported. One issue with conventional compensatory mutation analysis is that it requires both knowing an RNA’s function *a priori* and having a precise experimental assay for that function. Another issue is that lack of rescue does not provide information for or against the tested base pair; in general, several base pairs for each helix, mutated not only to their complement but also to other Watson/Crick pairs, need to be tested in these studies. Mutate-map-rescue (M^2^R) proposes to use chemical mapping as a general and high throughput readout of the experiment, even for RNAs whose functions are unknown or are difficult to assay (Figure 1C).

#### Mutate-map-rescue (M^2^R) results

Recent studies have established high-throughput mutate-map-rescue as a tool for rapidly validating or refuting RNA structure models, and have provided strong support for M^2^-derived models. For an *E. coli* 16S ribosomal RNA domain 126-235, modeling guided by 1D SHAPE data gave a solution-state secondary structure model different from the structure seen in the crystallized protein-bound small ribosomal subunit (Deigan et al., 2009). In contrast, M^2^ recovered a secondary structure that matched the crystallographic structure up to single-nucleotide register shifts, and M^2^R experiments involving 36 sets of compensatory mutations supported the M^2^ model, with no evidence for the one-dimensional SHAPE-based alternative structure (Tian et al., 2014). This study demonstrated the use of M^2^R as a tool for disambiguating fine-scale uncertainties, including register shifts. Figure 3D shows an additional example to distinguish between two register shifts of a helix P5/alt-P5 in the lariat-capping GIR1 ribozyme (S.T., R.D., unpublished data). The restoration of the chemical profile of the wild type RNA from double mutations predicted to rescue P5, but not alt-P5, was visually apparent and confirmed by the subsequently released crystal structure of the ribozyme (Meyer et al., 2014).

It is important to note here that the confident interpretation of M^2^R measurements does not require the ‘punctate’ release of partners nucleotides upon single mutations. For example, if single mutations of both partners in a base pair lead to alternative secondary structures with dramatically different chemical profiles [see Section 5, and several examples in (Tian et al., 2014)], M^2^ analysis would not provide clean evidence of their pairing. However, in M^2^R, restoration of the wild type profile upon double mutation would still provide strong experimental evidence for the base pairing of the nucleotides.

The M^2^R experiment has further provided strong tests of several stems of a recently discovered internal ribosome entry site (IRES) in the HoxA9 mRNA, including a previously uncertain pseudoknot predicted with low bootstrap support (56%) (Xue et al., 2015). Further cellular assays tested the *in vivo* relevance of the M^2^-rescue structural model, again through compensatory rescue but with a functional readout of IRES activity.

#### Prospects for higher-dimensional chemical mapping (mutate-mutate-map, M^3^)

The nucleotides targeted by M^2^R have been limited to base pairs that remain uncertain after M^2^ analysis. The method might, in principle, be generalized to cases in which no secondary structure hypotheses or energetic models are assumed or modeled *a priori,* as was the original goal of M^2^ (Section 3.2). Such a ‘model-free’ method would involve profiling the effects of *all* double mutants of target RNA on the chemical reactivities of all other nucleotides, and cataloging the pairs of mutations that rescue perturbations of single mutations. These data would give a ‘three-dimensional’ data set (Figure 1C); we refer to the procedure as a mutate-mutate-map analysis (M^3^). The expected sequencing costs of M^3^ (see Section 6 below) have prevented broad testing of the concept, although massively parallel synthesis and sequencing methods may allow such data sets to be collected for short transcripts. At present, the M^2^R method, which provides a targeted subset of a full M^3^ data set (Figure 1C), has turned out to be sufficient – and, in some cases, necessary – to achieve confidence in secondary structure models.

### 3.5 Acceleration from mutational profiling (MaP)

M^2^ measurements require separate synthesis and purification of single mutants of the target RNA. This is possible for RNA molecules that can be transcribed from DNA templates that can in turn be constructed through PCR assembly of small primers. This synthesis process is straightforward for domains up to a few hundred nucleotides but becomes difficult for RNAs of longer length or for transcripts that require *in vivo* biogenesis to assemble into functional structures. A method that yields M^2^-like data without single mutant libraries has recently been achieved (Homan et al., 2014; Siegfried et al., 2014). In this method, the initial perturbation to the RNA structure is not a mutation at an initially protected nucleotide but a chemical modification at that nucleotide when it is transiently available for modification. The effect of this first perturbation then affects the chemical modifications at other nucleotides that occur later in the reaction period (Figure 4A). Unlike conventional chemical mapping approaches where one typically seeks ‘single-hit’ modification kinetics (fewer than one average number of modifications per transcript), this protocol explicitly seeks multiple hits per transcript to enable detection of correlations between modification events at different sites. Detection of multiple hits per transcript was enabled by the development of <u>m</u>ut<u>a</u>tional <u>p</u>rofiling (MaP), a protocol for primer extension and next-generation sequencing that allows reverse transcriptases to bypass modification sites and incorporate mutations into the cDNA transcript instead of terminating at those sites (Siegfried et al., 2014).

**Figure 4.**
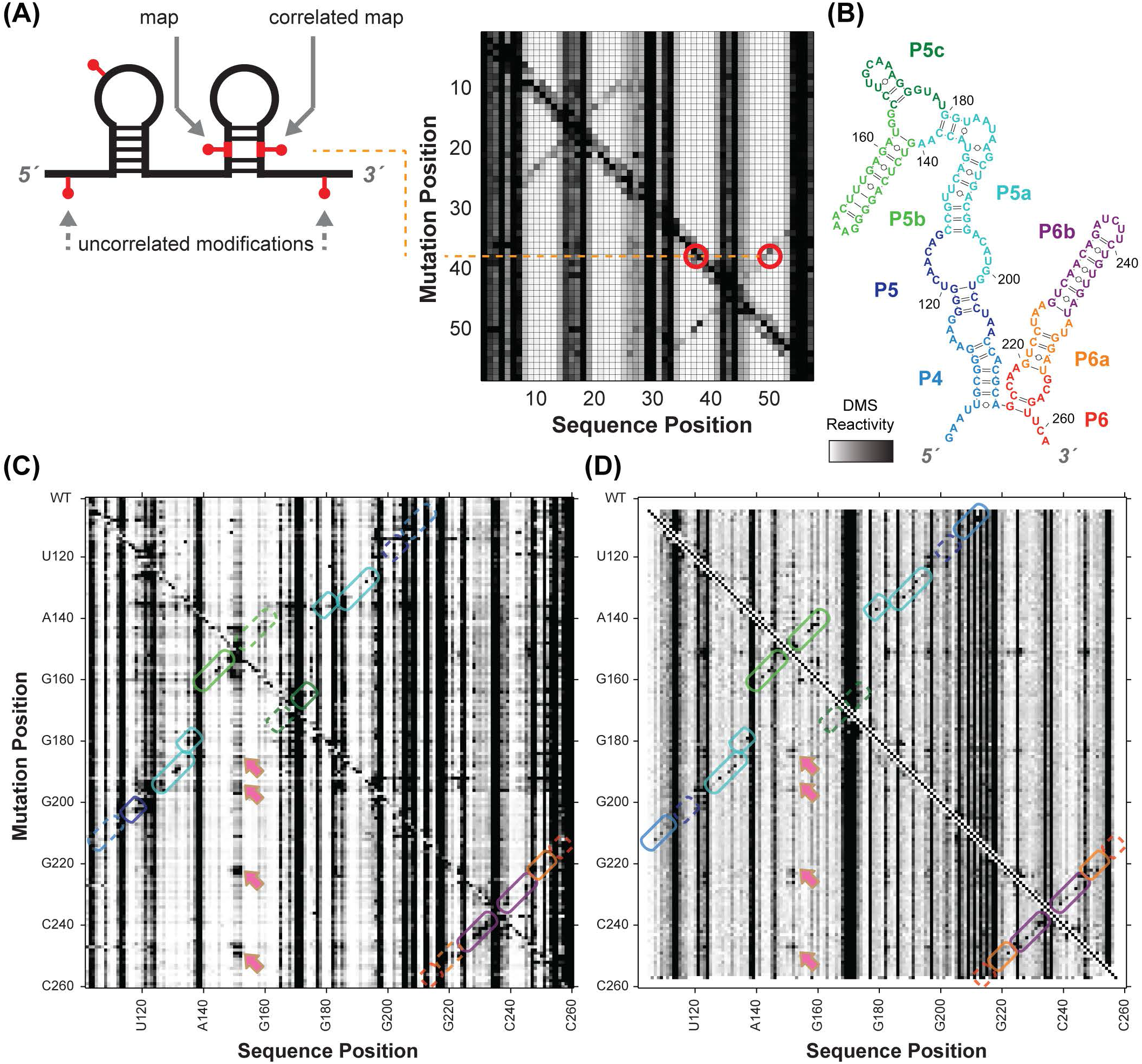
Schematic of single-molecule correlated modification mapping and data comparison for the *Tetrahymena* group I intron P4-P6 domain. (A). Schematic of how multiple modifications can read out RNA structure. A primary modification serves as a ‘mutation’ similar to M^2^, leading to a correlated secondary modification at its base-pairing partner. Multiple chemical modification events on the same RNA are read out by reverse transcription under conditions in which mismatch nucleotides are incorporated into cDNA at modification sites. Simulated data are shown. (B). Secondary structure of the*Tetrahymena* group I intron P4-P6 domain. (C). M^2^-DMS measurements for the P4-P6 RNA; helix features color-coded as in (B). (D). Data using DMS in multiple-hit conditions, collected previously for RNA Interaction Group (RING-MaP) analysis (Homan et al., 2014) but displayed here in a distinct ‘MaP-2D’ view. The rate of modifications at each nucleotide position, given a detection of nucleotide modification at every other position, is shown. Each row shows such a profile, normalized by the sum of counts at each position. In panels (C) and (D), red arrows mark exposure of the P5b loop upon disruption of the RNA tertiary structure from not only mutation of this loop’s ‘receptor’ (J6a/b) but also other helix perturbations. RMDB Accession IDs for datasets shown: (C). TRP4P6_DMS_0002; (D). adapted from (Homan et al., 2014).

For several RNAs, novel RING-MaP (<u>R</u>NA <u>in</u>teraction <u>G</u>roups by <u>M</u>utational Profiling) analysis of multiple-hit DMS data revealed statistically significant modification-modification correlations between several nucleotide pairs in the same helices, pairs involved in tertiary contacts, and pairs that were not directly in contact but might be exposed concomitantly in weakly populated states (Homan et al., 2014). Figure 4B shows an alternative two-dimensional view of these same data for the P4-P6 domain of the *Tetrahymena* ribozyme: a heat-map of the modification frequency at one site given that a modification is observed at a second site (S.T., R.D., C.Y.Cheng, unpublished data). This view, termed herein ‘MaP-2D’ analysis, illustrates the similarities between this protocol that maps correlations between multiple chemical modifications and the M^2^ approach (Figure 4C). In both panels, vertical striations correspond to the general one-dimensional DMS modification pattern: there is a high rate of modification at unpaired regions independent of where other modifications appear. Both panels also show detailed 2D information correlating the exposure of generally protected nucleotides with modifications at other nucleotides. Cross-diagonal features corresponding to all the RNA’s helices are visible as punctate dots (in colored outlines) as well as signals for the tetraloop/tetraloop-receptor tertiary contact (magenta arrows). Interestingly, in the MaP-2D data, a punctate signal at, for example, an A-U Watson-Crick base pair involves DMS modification at both the adenosine and a ‘non-canonical’ modification at the uracil. It is not yet clear if the latter events are due to modification at uracil transiently deprotonated at the N1 position or to other kinds of modification.

Given the visual similarity of the M^2^ and MaP-2D data, automated secondary structure analysis developed for M^2^ measurements apply readily to MaP-2D data, allowing the recovery of all helices of this as well as other RNA domains that have been challenging for chemical-mapping-derived secondary structure modeling (S.T., R.D., unpublished data). These results suggest that MaP-2D will be able to achieve data and secondary structure models with quality comparable to M^2^ through a simpler protocol that obviates preparation of sequence mutants. Independent validation procedures for MaP-2D experiments have not been developed, so testing the resulting models will still likely require synthesizing variants with single and double mutations and testing for compensatory rescue, as described in Section 3.4 above.

### 3.6 Summary

Critical benchmarks and blind tests of the mutate-and-map (M^2^) concept, high-throughput mutate-map-rescue, and MaP-2D have been carried out on more than a dozen RNA systems. These studies have supported the basic MCM hypothesis, especially with regards to secondary structure: multidimensional expansions of chemical mapping give rapid, automated, and consistently accurate solution-state structure models of RNA molecules.

## 4. Multiplexed ·OH cleavage analysis for 3D structure

### 4.1 MOHCA proof-of-concept

Many RNAs are known to form specific tertiary structures to carry out catalysis or to recognize small molecule, protein, or nucleic acid binding partners. While the studies above have supported application of M^2^ and related methods to infer secondary structure, these data have not in general returned information needed to resolve the global tertiary arrangement of those helices, much less atomic-resolution tertiary structure. Tertiary information from M^2^ has been limited typically to pseudoknots or a fraction of the structure’s other non-canonical base pairs, as in the P2/P9 A-minor interaction in the GIR1 ribozyme (Figure 3B). As an illustration of the difficulty of inferring non-canonical pairs, mutations in each A-minor interaction interconnecting the two aptamers of a double-glycine riboswitch successfully disrupted these interactions but also disrupted numerous other tertiary interactions as well (Kladwang et al., 2011b) (yellow arrows, Figure 3A). 3D modeling is difficult without such precise tertiary contact information, and has been carried out only for favorable cases such as an adenine riboswitch aptamer (Kladwang et al., 2011b) or at low resolution (Homan et al., 2014). Recent RNA-Puzzle blind trials further illustrate the problem: M^2^-guided 3D models with the correct global tertiary structure modeled at sub-helical (better than 1 nm) resolution have been submitted for most problems, but modelers have not been able to rank their most accurate submissions as their top models (Miao et al., 2015).

#### Precedents for pairwise data from tethered radical cleavage

A different MCM protocol has been developed to help address the need for high-throughput RNA tertiary proximities, based on RNA-tethered radical sources. The protocol involves chemical attachment of iron chelates to single positions in the RNA backbone during or immediately after *in vitro* synthesis. After folding, hydroxyl radicals (·OH) are produced from these iron centers via the Fenton reaction, with Fe(II) being regenerated from Fe(III) by a reducing reagent such as ascorbate. The radicals attack nucleotides that are at distances of 15-30 Å to the radical source; oxidation of sugars can result in backbone cleavage (purple arrows leading to red lightning bolt, Figure 5A). While probing distance scales 2-5 fold longer in distance scale than the ∽6 Å separation of adjacent nucleotides, these data are expected to be powerful for constraining tertiary folds. (An analogy to smaller distance scales may be helpful: NMR approaches achieve near-atomic resolution on small macromolecules using rich sets of NOE-derived proximities between atom pairs separated by 3-5 Å, several fold longer than the 1 Å atomic length scale.) Indeed, classic work with sources tethered to single residues of transfer RNA, ribosomes, and other non-coding RNAs calibrated the relationship of RNA backbone cleavage with distance and established the utility of these data for nucleotide-resolution RNA and RNA-protein modeling; see, e.g., (Bergman et al., 2004; Culver & Noller, 2000; Han & Dervan, 1994; Lancaster et al., 2002). The accuracy of pairwise constraints from tethered radical source experiments has been further supported by comparison of these and other types of biochemical data on the ribosome to subsequently solved crystal structures (Sergiev et al., 2001; Whirl-Carrillo et al., 2002).

**Figure 5.**
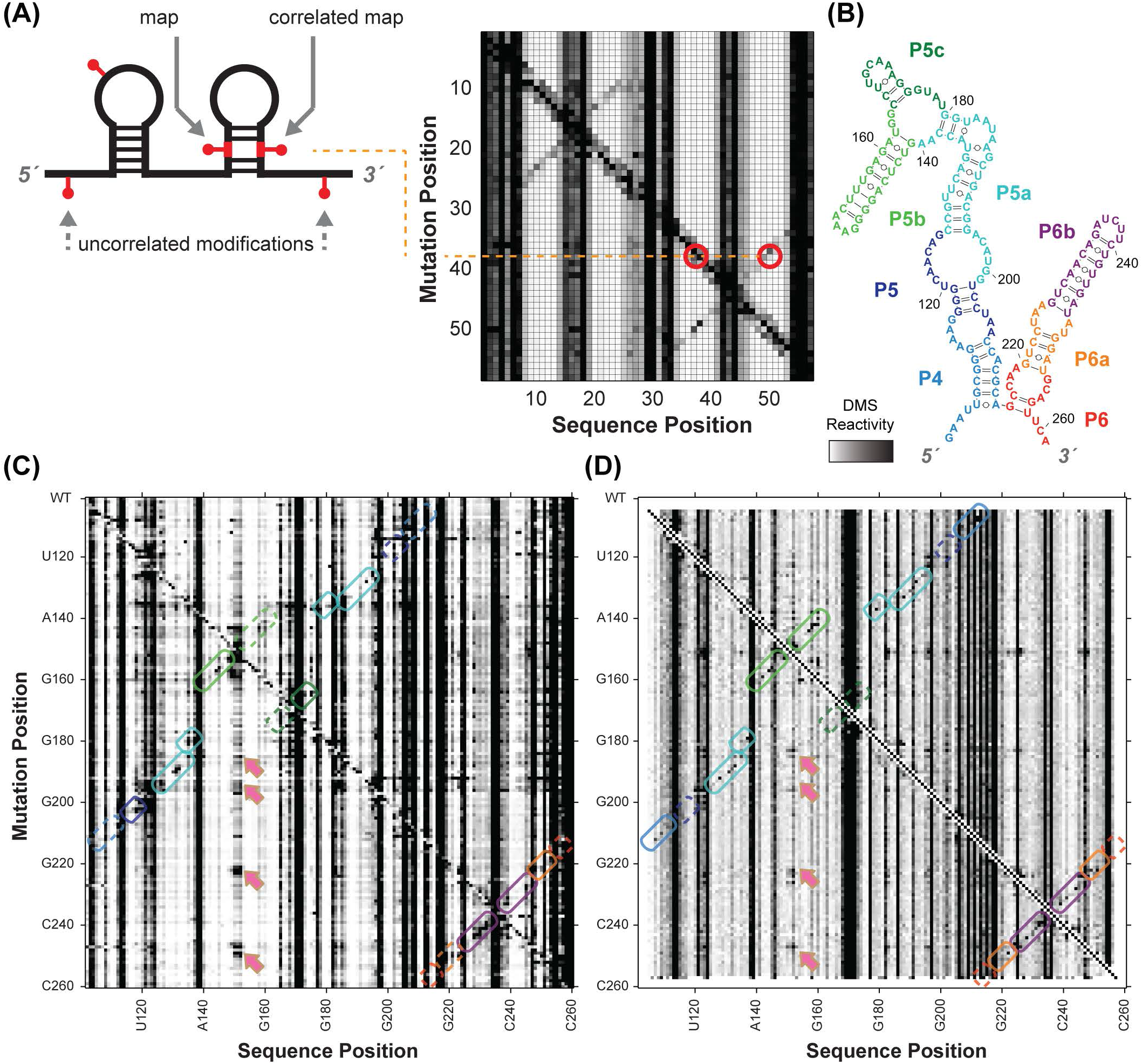
MOHCA-seq provides pairwise tertiary proximity information of RNA. (A). Schematic of MOHCA-seq (multiplexed OH cleavage analysis read out by deep sequencing). After generation of hydroxyl radicals (OH, purple), a strand scission event (red lightning bolt) and the corresponding iron chelate radical source position (yellow circle marked Fe) can be mapped out by subsequent reverse transcription to cDNA (green arrow) and paired-end sequencing. Simulated data are shown. (B). Additional oxidative damage events (red pins) that were not detectable in the original gel-based readout of MOHCA but are detectable by MOHCA-seq through termination of reverse transcription (green arrows). (C-E). MOHCA-seq data and tertiary structure models of (C) a double-aptamer glycine riboswitch from *F. nucleatum* with 10 mM glycine with cross-aptamer tertiary contacts (magenta arrows in MOHCA-seq map), (D) the GIR1 lariat-capping ribozyme from *D. iridis,* RNA-Puzzle 5, and (E) the ydaO cyclic-di-adenosine riboswitch with 10 μM c-di-AMP, RNA-Puzzle 12. The latter two are blind tests. Structures labeled ‘MCM predicted model’ were based on a multidimensional chemical mapping (MCM) pipeline of M^2^ secondary structure analysis, MOHCA-seq tertiary proximity mapping, and Rosetta computational modeling. Crystal structures are from the PDB, (C) 3P49, (D) 4P8Z, (E) 4QK8. In (D), red asterisks mark two positions that undergo catalytic modification (lariat formation and hydrolytic scission) by the ribozyme; for visual clarity, data at those positions are not shown. MOHCA-seq maps of (C-E) are filtered to show features with signal-to-noise ratios above 2 (different from a cutoff of 1 in (Cheng et al., 2015b)). Cyan contours highlight map features corresponding to each secondary structure helix. Other contours mark hits that were inferred through visual inspection of MOHCA-seq maps; to aid visual comparison, only contours including at least one residue pair with phosphorus-phosphorus (P-P) distance < 45 Å in the crystal structure are shown. Coloring of these tertiary contours reflect P-P distances of closest approach for residue pairs in the MCM predicted models (green, < 30 Å; yellow, 30 - 45 Å; red, > 45 Å). The same coloring is shown for cylinders in bottom panels of structures, which connect pairs of residues of closest distance corresponding to each contour; thick and thin cylinders correspond to strong and weak hits in (Cheng et al., 2015b). Each 3D model is shown with colored cylinders, or helices with matching color as in Figure 3. MOHCA-seq maps have colored axes matching secondary structure in Figure 3. In (E), gray spheres show position of two c-di-AMP ligands in both model and crystal structure. RMDB Accession IDs for datasets shown: (C). GLYCFN_MCA_0002; (D). RNAPZ5_MCA_0001; (E-F). RNAPZ12_MCA_0000.

#### MOHCA with gel readout

<u>M</u>ultiplexed •<u>OH</u> <u>c</u>leavage <u>a</u>nalysis (MOHCA) was reported in 2008 to give secondary and tertiary structure information on RNA structure from a chemical mapping method (Das et al., 2008). MOHCA involved random incorporation of radical sources at all possible sites of an RNA, identification of the positions of radical cleavage through gel electrophoresis, and identification of which source position produced which cleavage events through in-gel RNA scission at radical source sites and electrophoresis in a perpendicular direction. Data from this first MCM technique gave two-dimensional maps that reflect not base pairing, as in M^2^, but spatial proximity extending over tens of Angstroms. While necessarily lower in resolution, these maps confirmed the secondary structure of this RNA in several solution conditions and, crucially, described lower resolution proximities between helical elements arranged in space. MOHCA maps were sufficiently information-rich to guide Rosetta 3D modeling methods to a 13 Å-RMSD accuracy model of the tertiary structure of an RNA model system, the P4-P6 domain of the *Tetrahymena* ribozyme. The MOHCA-Rosetta method also gave initial ensemble models of the conformationally heterogeneous states of the P4-P6 RNA without magnesium. Several groups developed methods to incorporate MOHCA data into 3D computational methods (Jeon et al., 2013; Parisien & Major, 2012; Seetin & Mathews, 2011). However, the MOHCA experimental protocol required custom-synthesized nucleotides with double modifications (2’-NH**2** for source attachment; α-phosphorothioate for iodine-catalyzed scission), two-dimensional gel electrophoresis, and numerous gel replicates for separate 5’ and 3’ end-labeled samples and with different running times to resolve different lengths. These requirements prevented MOHCA from being subjected to blind tests or entering routine use for RNA structure inference.

### 4.2 Acceleration through MOHCA-seq

The advent of paired-end next generation sequencing resolved the difficulties of the original MOHCA method. An updated MOHCA-seq protocol has been developed, which uses commercially available nucleotides and iron chelate reagents to prepare the library of RNAs with radical sources (Cheng et al., 2015b). After folding and fragmentation, an RNA-seq-inspired protocol allows readout of radical cleavage events and associated source locations. Primer binding sites are ligated onto the cleaved RNA ends, and reverse transcription from these primers (green arrows, Figure 5A) terminate at the radical source. Unlike the original scission-based protocol, the reverse transcription can also terminate at and read out other oxidative damage events associated with the radical source, giving additional pairs of nucleotides that are both proximal to the radical source (red pins and lightning bolts, Figure 5B). A second adapter ligation step enables paired-end sequencing of these cDNA fragments and determination of these pairs of nucleotides. Because the final data are digital, background subtraction, correction for reverse transcription attenuation, and error estimates can be carried out through an automated procedure (closure-based •OH correlation analysis, COHCOA). Single MOHCA-seq experiments give data as rich as experiments involving dozens of gels with the original MOHCA method, mainly due to the readout of double-modification events (Figure 4B) and the ability to carry out digital data processing.

In a benchmark on five RNA domains of known structure with lengths up to 188 nucleotides, MOHCA-seq maps consistently gave signals that confirmed the RNA’s solution-state secondary structure and, most importantly, gave information that enabled tertiary structure modeling. For a double glycine riboswitch aptamer, all six helices observed previously with M^2^ (Figure 3A) gave distinct hits in MOHCA-seq data (black features inside cyan contours, Figure 5C). Furthermore, the MOHCA-seq map marked riboswitch regions brought together by cross-domain A-minor contacts (magenta arrows in Figure 5C), information that could not be resolved by M^2^ (Figure 3A) due to cooperative loss of all cross-domain tertiary contacts upon mutation.

While these interactions could be seen through visual inspection, the MOHCA-seq map did not allow compilation of a complete list of non-canonical pairs at nucleotide resolution, much less a global tertiary structure model. On the tested domains and in prior work (Cheng et al., 2015b; Das et al., 2008; Sergiev et al., 2001; Whirl-Carrillo et al., 2002), the median distance of MOHCA-seq-connected hits was ∽30 Å, on the same scale as the diameter of an RNA helix (26 Å) and larger than the ∽6 Å sugar-to-sugar separation of sequence-adjacent nucleotides. This intrinsic resolution is unlikely to improve significantly, even if the iron-chelate can be tethered more closely to the RNA, since pairs of nucleotides that are brought into distance much closer than 15 Å are typically buried within contacts and protected from radical attack. Given this likely intrinsic limit in resolution, achieving 3D structural pictures requires integration of MOHCA-seq data with *de novo* computational methods, analogous to the integration of M^2^ analysis with automated algorithms to give secondary structures (Section 2.3).

### 4.3 Tests for MCM 3D modeling

#### Integration with computational tertiary structure modeling

To test its information content for 3D structure, MOHCA-seq was integrated with the Rosetta <u>F</u>ragment <u>A</u>ssembly of <u>R</u>NA with <u>F</u>ull <u>A</u>tom <u>R</u>efinement (FARFAR) method for 3D structure modeling (Cheng et al., 2015a). Analogous to the guidance of RNA secondary structure prediction with M^2^ data (Section 3.2), a list of nucleotide pairs with strong MOHCA intensities was compiled for each RNA. A low-resolution scoring function guides initial FARFAR modeling, and 3D structures that brought these pairs of nucleotides were awarded an energy/score bonus. When carried out using the benchmark data described above and taking advantage of M^2^ data to predefine secondary structure, this M^2^-MOHCA-Rosetta pipeline achieved 8-12 Å RMSD accuracies, a resolution that allowed accurate visualization of the tertiary arrangement of helices at near-nucleotide resolution (Figure 5C). Modeling without MOHCA-seq data gave significantly worse RMSD (e.g., 30.5 Å instead of 7.9 Å for the glycine riboswitch aptamer), confirming the necessity of these MCM data. For a newly discovered HoxA9 mRNA IRES, MOHCA-seq data supported a secondary structure and pseudoknot detected by previous M^2^R experiments (Xue et al., 2015) and allowed 3D modeling of the RNA as a ‘loose tertiary globule’ (Cheng et al., 2015a).

#### Blind tests

As with the secondary structure tests for M^2^, the most important tests of MOHCA-seq tertiary structure inference have been blind trials. To date, two partial blind tests have been carried out. The first involved refinement of nearly 40% of a GIR1 lariat-capping ribozyme model before the release of this RNA-Puzzle’s crystal structure (Figure 5D). The MOHCA-seq-guided refinement indeed improved the accuracy of the refined regions from 17.0 Å to 11.2 Å and, for the whole ribozyme, from 9.6 Å to 8.2 Å (Cheng et al., 2015b). A second blind test involved an RNA-Puzzle on a cyclic-di-adenosine monophosphate riboswitch aptamer (Ren & Patel, 2014). In this case, the MOHCA-seq protocol (which had only recently been developed) was carried out on the target molecule only a few days before the modeling deadline, too late to influence modeling. Nevertheless, *post facto* comparisons highlighted the discriminatory potential of MOHCA-seq maps. Several MOHCA-seq hits involved residue pairs that were more than 45 Å distant in the submitted models (MCM Predicted Model, Figure 5E), but these discrepancies were resolved when plotting distances derived from the subsequently released crystal structure (Crystal Structure, Figure 5E). These results suggest that inclusion of MOHCA-seq data during 3D modeling could significantly improve accuracy. Collection and dissemination of MOHCA-seq data for more recent RNA-Puzzles are offering further rigorous tests of this hypothesis (C.Y. Cheng, M. Magnus, K. Kappel, R.D., unpublished data).

### 4.4 Towards mature MOHCA-seq modeling

The above studies have given initial support to the overall hypothesis that MOHCA-seq can complement M^2^ to produce RNA 3D models with useful sub-helical resolution. Nevertheless, there are at least two important aspects of the tertiary structure modeling that are underdeveloped in comparison with the M^2^-based secondary structure modeling: uncertainty estimation and independent validation protocols.

First, the studies above gave estimates of the 3D modeling precision based on the similarity of different low energy models from a single computational modeling run, but these values may be biased towards overestimating accuracy, as occurs in NMR modeling (Rieping et al., 2005). A bootstrapping procedure, similar to that used for M^2^-derived secondary structure models in Section 3, might achieve more conservative estimates. While resampling MOHCA-seq constraint lists can already generate bootstrapped ‘mock’ data sets, Rosetta modeling is currently too computationally expensive to allow replicate runs with these data sets. Accelerations in Rosetta modeling, or use of alternative 3D modeling protocols (Krokhotin et al., 2015; Parisien & Major, 2012), will be needed to attain such uncertainty estimates.

Second, there is no tertiary structure analog yet of the compensatory rescue experiments that test secondary structure. MOHCA-seq modeling does not typically resolve individual base pairs of RNA tertiary contacts, precluding design of compensatory mutations. Even the modeling could achieve such resolution, most tertiary interactions involve non-canonical pairs, often making additional interactions with other nucleotides. These pairs are not expected to be replaceable with alternative pairs without energetic cost.

As an alternative, one can envision a motif-level testing procedure involving substitution of entire motifs of the RNA. For example, if a 3D MCM model predicts a sharp bend and twist at a two-way junction, one could replace that junction with a previously solved junction known to form a similar bend and twist. Positive evidence for the predicted junction geometry would come from chemical mapping or functional experiments showing that separately substituting one strand or the other produces a disruption in 3D structure/function and that concomitant mutation rescues the structure/function. Similar replacements for three-way, four-way, and higher order junctions and for tertiary contacts might also be feasible. One challenge for this motif-by-motif approach would be to automatically find and design the appropriate substitutes. It is presently unclear if the database of known structures is large enough to provide such substitutes. Another challenge would be to ensure that false positives do not arise from simple rescue of secondary structure rather than tertiary structure. The development of incisive testing procedures of 3D model features, analogous to compensatory rescue of Watson/Crick pairs, is an important frontier for MCM and other RNA structural biology methods, especially as they seek to visualize transcripts whose functionally relevant structures may only form *in vivo.*

### 4.5 Summary

Benchmarks and a blind test of the MOHCA (multiplexed ·OH cleavage analysis) concept for RNA proximity mapping have been carried out on nearly a dozen RNA systems. Complementary to mutate-and-map data that pinpoint RNA secondary structure, MOHCA seeks proximal nucleotide pairs that would enable computer modeling of RNA tertiary structure at nanometer resolution. The studies to date have extended support of the basic MCM hypothesis from secondary to tertiary structure: multidimensional expansions of chemical mapping enable consistently accurate three-dimensional structure models of RNA molecules.

## 5. Deconvolving multiple RNA structures with MCM

### 5.1 Multiple states of RNA as a major challenge

As noted in the Introduction, most biological RNA molecules that have been studied in detail transit through multiple structures during their functional cycles. For example, viral RNA genomes interconvert between compact structures in packaged forms, less-structured cellular states that can recruit and organize host proteins, and states available for translation or replication [see, e.g., (Bothe et al., 2011; Filbin & Kieft, 2009; Schneemann, 2006) and refs. therein]. On one hand, one-dimensional chemical mapping data are sensitive to multiple structures, and recent studies *in vivo* and *in vitro* support a picture of many, and perhaps most, regions of RNA transcripts interconverting between complex conformational states [see, e.g., (Kwok et al., 2015; Rouskin et al., 2013; Spitale et al., 2015)]. On the other hand, whether these conformational changes are functional or simply ‘structural noise’ is unknown for most regions, and the uncertainty is exacerbated by the difficulty of deconvolving the component structures from data that average over the entire ensemble of structures (Eddy, 2014; Washietl et al., 2012). Multidimensional chemical mapping measurements give rich data on RNA structure and, in favorable cases, allow deconvolution of ensembles of secondary and tertiary structures from experiments.

### 5.2 Deconvolving riboswitch secondary structures with M^2^-REEFFIT

Although M^2^ measurements were not originally developed to deconvolve multiple states of an RNA, early measurements suggested that these data captured evidence of alternative states. Even for well-structured RNAs, some single mutations produce changes in chemical reactivity over extended regions (yellow arrows in Figure 3A-3C), and similar patterns of changes occur in several mutants. The secondary structure dominating the RNA ensemble apparently shifts to a distinct secondary structure in those variants. Indeed, for certain RNAs, the majority of mutations have been observed to produce such delocalized rearrangements. Examples have included riboswitches that are known from other techniques to form multiple structures, engineered sequences that failed to fold into target structures, and engineered switches explicitly designed to form multiple structures (Figure 6) (Cordero & Das, 2015; Lee et al., 2014; Reining et al., 2013; Serganov et al., 2004). For these cases, it is not possible to define a single secondary structure for the RNA, and a separate analysis method has been developed that models an ensemble of secondary structures and, importantly, estimates the associated increase in modeling uncertainty.

**Figure 6.**
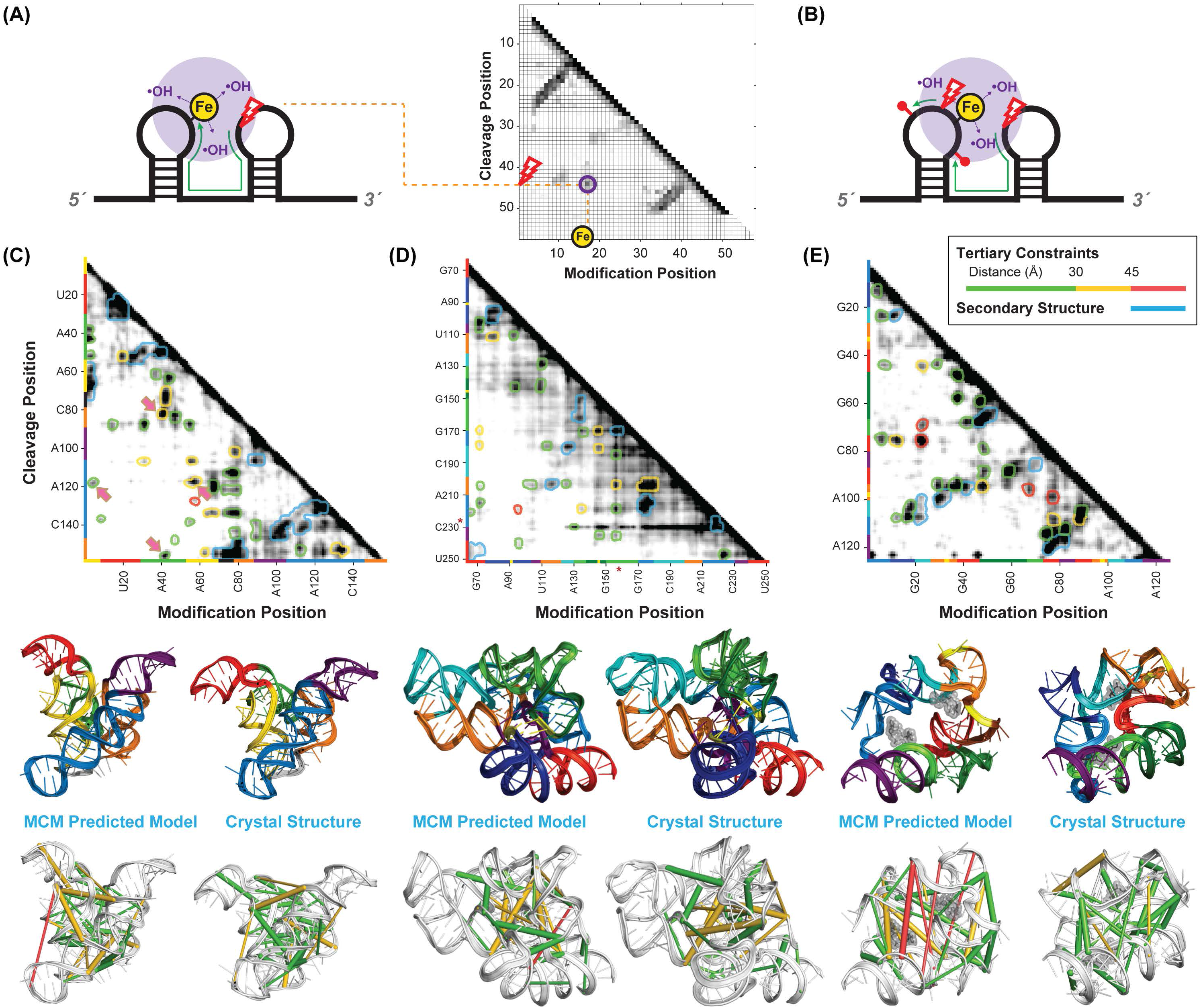
M^2^-REEFFIT reveals hidden states in secondary structure ensembles. (A, B). M^2^ data (left), fitted cluster weights (center), and fits from <u>R</u>NA <u>e</u>nsemble <u>e</u>xtraction <u>f</u>rom footprinting insights technique (REEFFIT, right) of the ‘Tebowned’ riboswitch designed to interconvert between two states upon binding of flavin mononucleotide (FMN). RNA was probed (A) in absence of FMN and (B) in presence of 2 mM FMN. Red rectangles in (A) mark nucleotide A30, which was not expected to be reactive in either of two target states of the riboswitch, but is explained by a third state uncovered by REEFFIT. (C). Secondary structures of REEFFIT predicted states. TBWN-A and TBWN-B were target states of the riboswitch design problem; TBWN-C was an unexpected state modeled by REEFFIT. (D) Prospective tests of REEFFIT model. 1D-SHAPE profiles of each state-stabilizing mutant agree well with the SHAPE profiles predicted from REEFFIT analysis. Red rectangle marks nucleotide A30, predicted and confirmed to be exposed in TBWN-C-stabilizing mutants. Data are from (Cordero & Das, 2015). RMDB Accession IDs for datasets shown: (A). TBWN_1 M7_0000; (B). TBWN_1M7_0001; (C). TBWN_STB_0000.

Modeling of full conformational ensembles from experimental data is a general problem in structural biology that is under active investigation in many labs. Since data must be used to infer not just a single structural model but instead the weights of a potentially large number of structures, no experimental method can directly read out an ensemble in a ‘model-free’ manner. Several approaches being currently developed for ensemble modeling find the minimal perturbations to a predefined, physically reasonable ensemble model that are necessary to recover experimental observables [see, e.g., (Beauchamp et al., 2014; Pitera & Chodera, 2012; Stelzer et al., 2011; van den Bedem & Fraser, 2015) and refs. therein]. <u>R</u>NA <u>E</u>nsemble <u>E</u>xtraction from <u>F</u>ootprinting <u>I</u>nsights <u>T</u>echnique (REEFFIT) is the first such approach developed for M^2^ data (Cordero & Das, 2015). The prior ensemble comes from automated prediction of equilibrium secondary structure ensembles. REEFFIT assumes that M^2^ data reflect a mixture of RNA secondary structures whose relative populations are shifted with mutation. While similar in concept to spectral analysis or principal component methods (Halabi et al., 2009; Homan et al., 2014), REEFFIT provides detailed models of the full ensemble and can make additional predictions. The method optimizes the ensemble model’s posterior probability, based on a well-defined likelihood model and Bayesian priors defined by empirical relationships between RNA pairing and chemical reactivity and the initial model of population fractions of each structure within each mutant, estimated from current RNA secondary structure energetic models. Figures 6A and 6B shows an example of M^2^-REEFFIT applied to understand an imperfectly engineered switch.

The probed multi-state RNA was designed as part of the EteRNA internet-scale RNA engineering project, which seeks basic design rules for RNA structure and function (Lee et al., 2014). The molecule was designed to change its favored structure in response to flavin mononucleotide (FMN); chemical mapping confirmed the desired behavior for the starting sequence as well as for a large number of mutants. However, these data suggested that a region near nucleotide 30 that should have been protected prior to FMN binding was instead exposed (red rectangle, Figure 6A). In this case, automated REEFFIT analysis provided a satisfactory fit to the entire data set (Figures 6A-B, right panels), automatically recovering the desired two states (TBWN-A and TBWN-B, respectively; the state names derive from the sequence’s name ‘Tebowned’). As expected, the populations of these states (their ‘weights’ in the secondary structure ensemble) varied in different mutants (middle panels, Figures 6A and 6B), allowing automated estimation of the component reactivities, and the populations in the starting sequence were 56 ± 16% and 27 ± 12%. In addition, REEFFIT exposed an unexpected third state (TBWN-C, population 17 ± 11%, Figure 6C), which explained the anomalous reactivity of A30 (Figure 6D). Each states’ population in the starting sequence was greater than expected from the modeling uncertainty, estimated through bootstrapping, motivating further tests. As predicted, the population of TBWN-B, which presents an FMN aptamer sequence in the correct secondary structure context, increased significantly in conditions with FMN (compare weights in Figure 6B to Figure 6A). Additional evidence for the three states and modeled structures came from design of mutations to strongly stabilize each mutant (Figure 6D); when synthesized, these constructs gave chemical reactivity patterns in agreement with predictions from REEFFIT on the original M^2^ data (Cordero & Das, 2015).

#### Current limitations to secondary structure ensemble modeling

Applications of M^2^-REEFFIT to date have been limited to sequences of lengths of 100 nucleotides or less due to the computational expense of optimizing energies of structural ensembles. For longer RNAs, alternative structure detection methods that produce less detailed pairing information but require less computational power, such as RING-MaP, may be more appropriate tools for automatically detecting alternative secondary structure states in MCM data. Nevertheless, in any of these methods, the problem of validating proposed alternative structures remains a challenge. Unlike M^2^R in single-structure RNAs (Section 3.4), compensatory mutations that restore the stability of a weakly populated structure are expected to change the population of this state relative to others in the RNA’s ensemble, leading to chemical mapping profiles different from the starting sequence even if ‘rescue’ of the single target structure is successful. For the cases to date, isolation of predicted alternative structures through the design of multiple stabilizing mutations has provided evidence of those structures, but these experiments neither constrain the population of these states in the starting sequence nor reveal whether those populations might be biologically relevant. While an independent approach, MOHCA-seq, gave independent support to M^2^-based secondary structure models above (Figure 5), RNAs with multiple structures give diffuse, low signal-to-noise MOHCA-seq maps that do not strongly falsify or validate secondary structure ensembles (W. Kladwang, R.D., unpublished data). It seems likely that strong tests of MCM detections of alternative states will require probing their involvement in an RNA’s functional cycle. In cases where the function is known, compensatory mutation/rescue can be read out through functional assays interpreted in detailed kinetic and thermodynamic frameworks. Such studies have been carried out for RNA machines and viruses but require significant specialized effort; see, e.g., (Fica et al., 2013; Villordo et al., 2010).

### 5.3 Preformed tertiary contacts in heterogeneous states with MOHCA-seq

In addition to the alternative secondary structures probed by M^2^, information on 3D conformational dynamics can be captured by MCM data. RNAs like the ribosome and some aptamer domains of riboswitches have largely preformed secondary structure but transit through multiple states as they fold or transit through their functional cycles; for examples, see (Baird et al., 2010a; Baird et al., 2010b; Behrmann et al., 2015; Das et al., 2003; Noller, 2005). For most RNAs with this property, however, it is unclear if tertiary structures are retained throughout the conformational cycle. For example, one-dimensional chemical mapping measurements applied to riboswitch aptamers for glycine and for adenosylcobalamin, show loss of protections around ligand binding sites and in tertiary contacts in ligand-free states compared to ligand-bound states (Kwon & Strobel, 2008; Nahvi et al., 2004; Sudarsan et al., 2008), but these data do not resolve whether these tertiary structural features might still be present at low population in the ligand-free states. In contrast, MOHCA-seq positively detects tertiary interactions in these three aptamers even in ligand-free states. These hits occur at the same residue-pairs that give hits in ligand bound states, but at lower strength (Figure 7) (Cheng et al., 2015b). These measurements, along with mutational analysis, suggest that the contacts are sampled transiently and perhaps without well-defined base pairing; such contacts would otherwise be difficult to resolve without specialized experiments such as single molecule FRET studies with probes introduced at interacting residues. The MOHCA-seq data do not constrain whether these transient contacts occur independently or in an all-or-none fashion; ensembles based on low-energy conformations from MCM-guided Rosetta modeling (Figure 7) give initial visualizations but are, at present, difficult to test or refine. Future work involving systematic mutagenesis coupled to a MOHCA readout (’mutate-and-MOHCA’) and computational methods that produce better-converged 3D ensembles may enable the expansion of REEFFIT-like procedures for data-driven secondary structure ensemble modeling to tertiary structure ensemble modeling.

**Figure 7.**
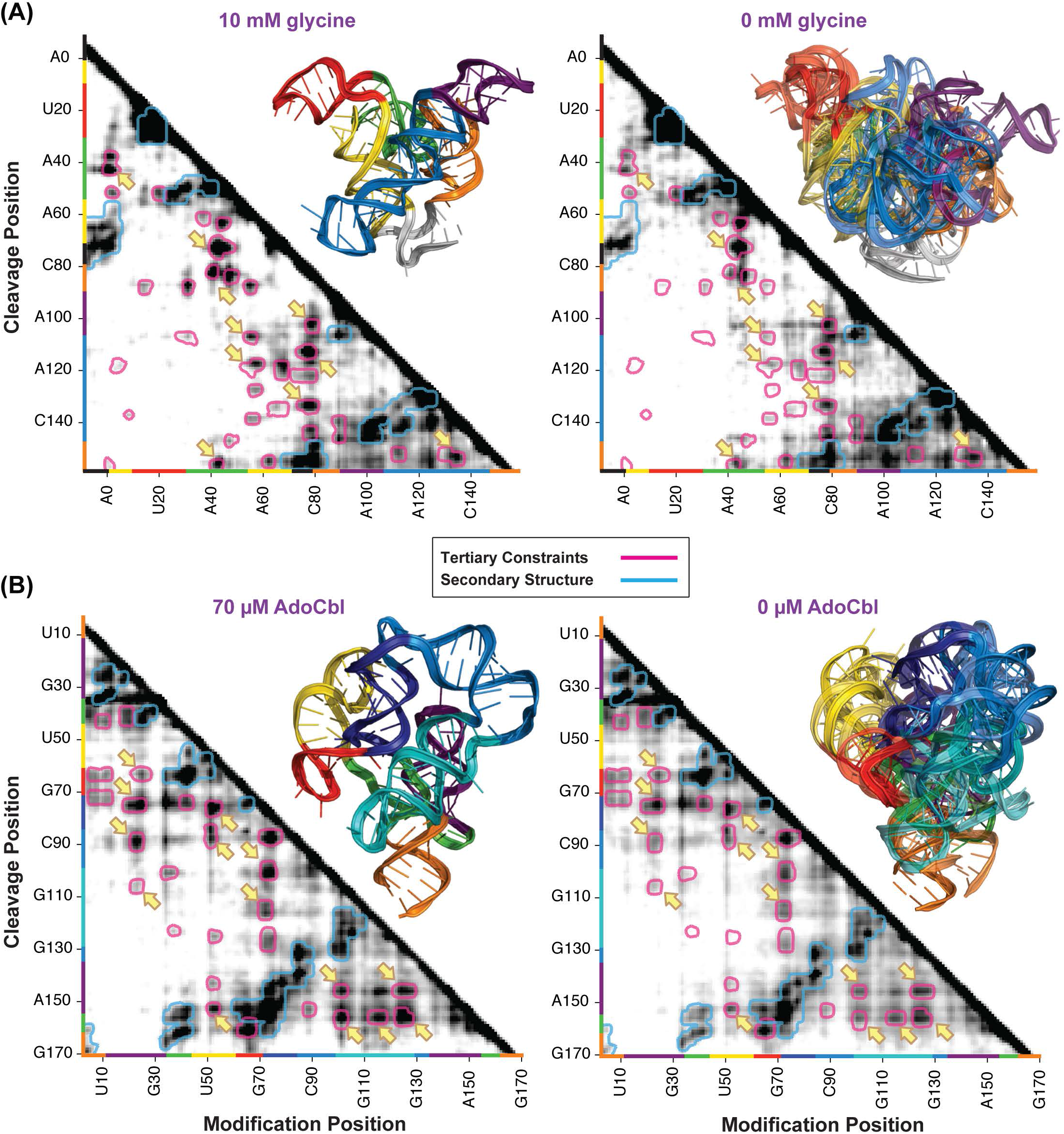
MOHCA-seq detects preformed tertiary contacts in riboswitches. MOHCA-seq data and tertiary structures (A) a double glycine riboswitch from *F. nucleatum.* RNA (including a kink-turning forming leader sequence), probed in presence of 10 mM glycine (left) or in absence of glycine (right); and (B) an adenosylcobalamin (AdoCbl) riboswitch from *S. thermophilus* (Peselis & Serganov, 2012), probed in presence of 70 μM AdoCbl (left) or in absence of AdoCbl (right). In each right panel, five MCM predicted models with lowest Rosetta energy provide an initial visualization of the ligand-free ensemble compared to the ligand-bound crystallographic structure (left panel). MOHCA-seq map filtering and color-coded contours in left panels (ligand-bound states) are same as in Figure 5, except that contours for tertiary contacts are colored uniformly in magenta. The same contours are shown in right-hand panels (ligand-free states). Yellow arrows point to regions in the MOHCA-seq maps showing tertiary contacts in the ligand-bound states (left) that appear at lower intensity in the ligand-free states (right). To avoid clutter, not all such hits are shown. RMDB Accession IDs for datasets shown: (A). CDIGMP_MCA_0002 and CDIGMP_MCA_0003; (B). GLYCFN_KNK_0005 and GLYCFN_KNK_0006; (C). RNAPZ6_MCA_0002 and RNAPZ6_MCA_0003.

### 5.4 Summary

Non-coding RNA states without single secondary or tertiary structures are functionally important and likely pervasive *in vivo* but available experimental methods have difficulty in characterizing them. Benchmarks of M^2^-REEFFIT support its use to recover known secondary structure ensembles and to detect unexpected alternative RNA structures. Extending MCM to flexible tertiary structures, application of MOHCA-seq to ligand-free states of three riboswitch aptamers detects preformed RNA tertiary contacts. These results support the use of multidimensional chemical mapping to visualize RNA states that involve heterogeneous secondary structure or tertiary structure.

## 6 Towards solving RNA structures *in vivo* with MCM

### 6.1 Upcoming challenges: from *in vitro* to *in vivo*

The development of multidimensional chemical mapping (MCM) techniques raises the prospect of *de novo* secondary structure and tertiary structure inference for the rapidly growing number of RNA molecules discovered in cells and viruses. Nevertheless, all MCM studies have been carried out *in vitro,* with separate experiments on each model system. Can MCM methods be extended to myriad RNA molecules interacting with their numerous other partners in their actual cellular or viral milieus? Several challenges will have to be solved before this is feasible.

#### Protection of RNA within RNPs and complexes

In terms of RNA biophysical states, it is possible that the binding of proteins and other partners will protect structurally important residue pairs from chemical modification and therefore obscure readout via MCM methods. Tests on the ribosome fully assembled with proteins and on riboswitches complexed to large ligands suggest that protections from molecular partners still leave significant MCM-detectable nucleotide-nucleotide pairing information [(Cheng et al., 2015b) and C.Y. Cheng, R.D., unpublished data]. However, the secondary structure and tertiary structure modeling methods that are currently used to integrate MCM data will need to take into account the possibility of these protections.

#### Making chemical perturbations and modifications *in vivo*

All MCM methods require perturbing and reading out structural effects on transcripts at single nucleotide resolution. In terms of chemistry, several methods are now available for making chemical modifications in cells and then reading out modification sites by high-throughput sequencing [reviewed in (Ding et al., 2014)]. It has also long been possible to make single-nucleotide-level perturbations on entire transcriptomes through, e.g., chemical mutagens. Correlating the perturbations with their structural effects may be possible with RING-MaP/MaP-2D-style protocols but will be challenging. More fundamentally, these modifications will generally disrupt more than just localized RNA structure. The modifications will lead to loss of functional interactions for each transcript, activation of stress responses, and other cell-wide perturbations, possibly including cell death. Possible solutions to this issue may be rapid delivery of chemical probes and quenching, as has been carried out in recent *in vivo* DMS and SHAPE measurements, although RING-MaP/MaP-2D approaches will require significantly higher modification rates than achieved in those experiments. Alternatively, MCM protocols might seek to introduce correlated chemical modifications into flash-frozen cells. For example, literature reports suggest that double-hit correlated modifications arise during irradiation of nucleic acids in frozen samples, although tests have only been described for double-stranded DNA (Chatterjee et al., 1994; Krisch et al., 1991).

#### Computational challenges

Obtaining structural models from MCM data requires integration via computational methods. Even for the least computationally expensive of these methods, which predict secondary structure without taking into account pseudoknots, modeling molecules longer than 2000 nucleotides remains challenging. For methods seeking three-dimensional structure at sub-helical resolution, molecules longer than 200 nucleotides have been intractable, even with predefined secondary structures and use of supercomputers, and, even for these cases, some steps remain non-automated (Cheng et al., 2015a; Miao et al., 2015). Multi-scale computational pipelines involving low resolution domain parsing, separate 3D folding of domains, assembly, and refinement will need to be developed as MCM data become available for viruses, ribosomes, and other large transcripts.

#### Sequencing costs

Multidimensional chemical mapping methods seek more information than one-dimensional chemical mapping approaches, and thus necessarily incur larger sequencing costs, e.g., in terms of necessary numbers of reads. Since MCM perturbs every nucleotide of an RNA and assays the response of every other nucleotide, the number of measurements should scale quadratically with the number of nucleotides (see, however, Section 6.2 below). Will these quadratic costs be acceptable for large RNA transcripts? To get a preliminary answer, we have estimated the minimal number of reads needed to achieve signal-to-noise acceptable for modeling secondary structure (M^2^, MaP-2D) or tertiary structure (MOHCA-seq), through sub-sampling from available data sets (see Figure 8 and its legend). As expected, the numbers of reads required to obtain such good quality data sets fit to a quadratic dependence with RNA length (Figure 9A). Compared to mutate-and-map (M^2^), which uses only the terminus of the read (orange), the MaP-2D analysis of RING-MaP data (red) gain efficiency by resolving multiple modification events via different mutations in a sequenced fragment. However, the most informative modifications occur at nucleotides that are most frequently sequestered into structure; these nucleotides contribute the fewest reads to the MaP-2D data, while they are mutated one-by-one in M^2^ to ensure sufficient signal-to-noise at all nucleotides. Overall, MaP-2D ends up requiring ∽50% more reads than M^2^. For tertiary contact discovery, MOHCA-seq maps are necessary, but those maps give comparably few features that report tertiary information compared to features that report secondary structure helices (compare number of cyan contours with all contours in Figure 5). Acquiring MOHCA-seq data for tertiary structure modeling is thus 2-3 times more expensive than M^2^ and MaP-2D data that target secondary structure (compare purple to red and orange curves, Figure 9A).

**Figure 8.**
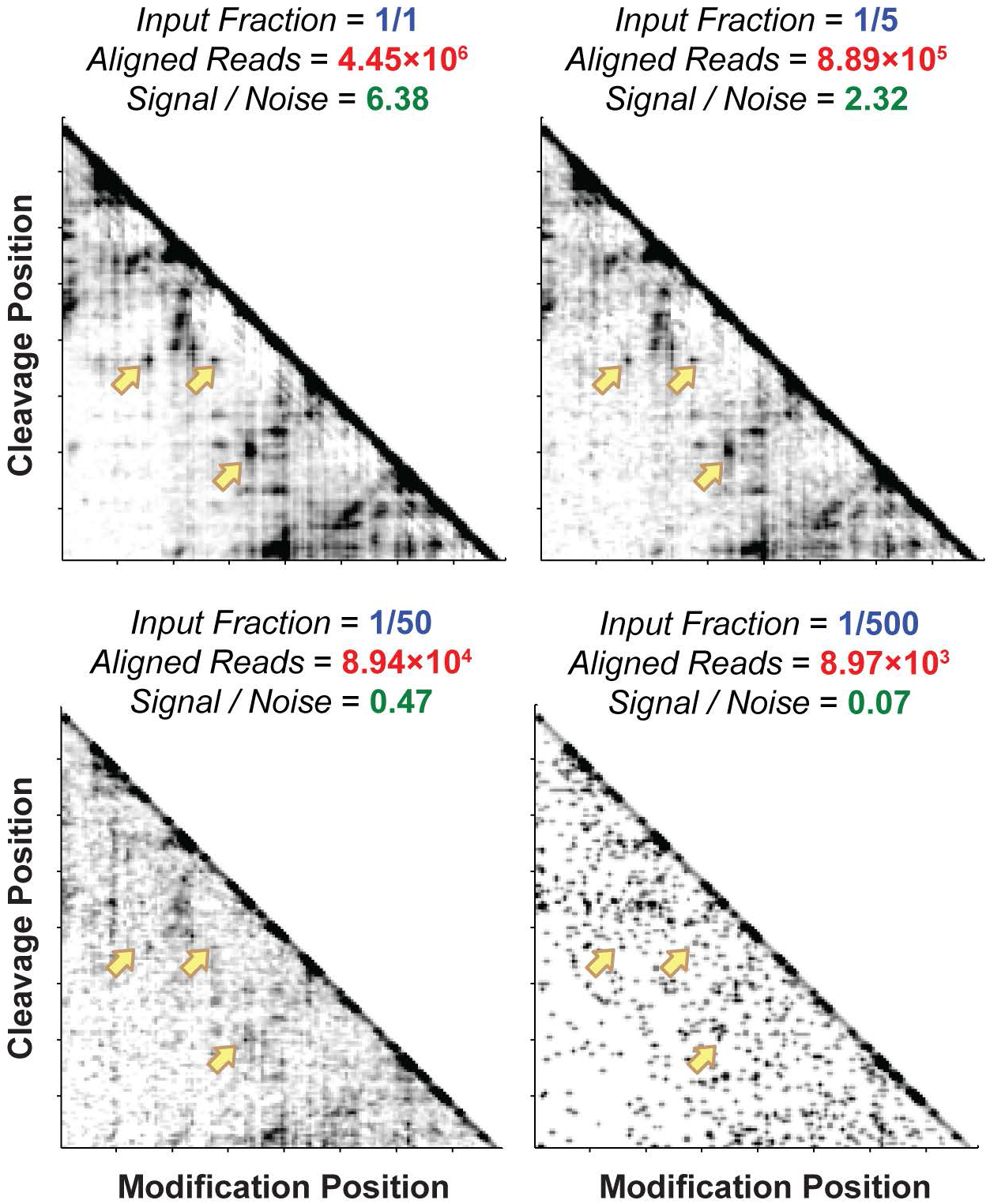
Subsampling of MCM data to determine minimal number of sequencing reads to infer RNA structure. MOHCA-seq data of a double glycine riboswitch from *F. nucleatum* were used (see also Figure 3A). A subset (1, 1/5, 1/500, and 1/5000) of the raw FASTQ file was randomly resampled and subjected through the complete COHCOA data processing and error estimation pipeline (Cheng et al., 2015b). Signal-to-noise ratio was estimated as the ratio between the mean of reactivity and the mean of error across the whole data set. Yellow arrows point to tertiary features that disappear as the number of resampled reads decreases. RMDB Accession IDs for datasets shown: GLYCFN_MCA_0002.

**Figure 9.**
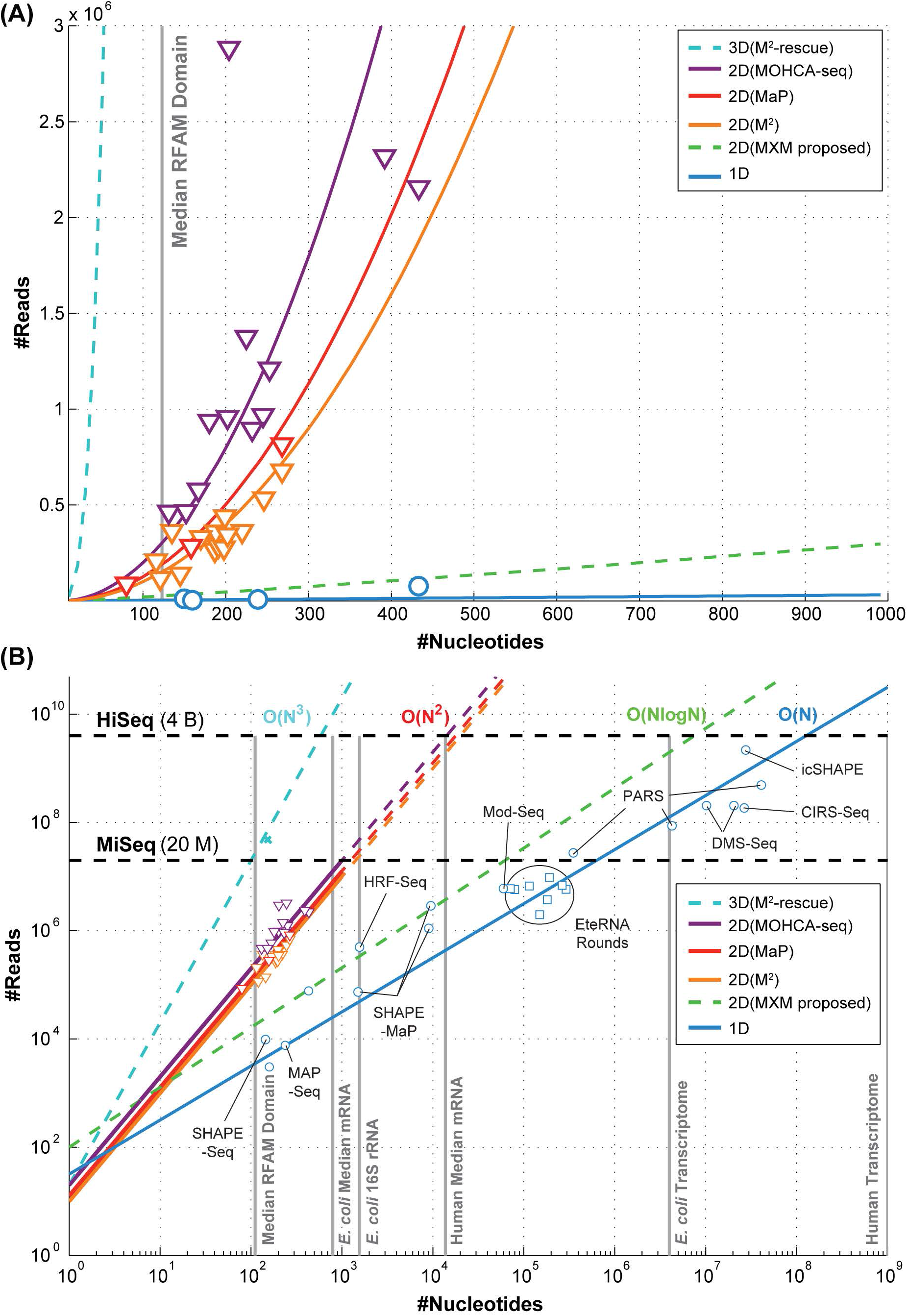
Scaling of sequencing costs for MCM. Expected sequencing costs (number of reads) versus RNA lengths, plotted on (A) linear scale and (B) logarithimic scale, required to achieve usable signal-to-noise levels for for 1D, 2D, and 3D chemical mapping methods described or proposed in text. Costs are estimated based on publicly available data for a number of RNAs and transcriptomes and the subsampling procedure described in Figure 8. Most M^2^ data (orange triangles) were collected by capillary electrophoresis (CE); conversion to number of Illumina reads was achieved by comparison of signal-to-noise values of CE and Illumina data sets for a 16S rRNA 126-235 four-way junction, for which both measurements are available. References for next-generation sequencing technologies for 1D mapping (blue circles): SHAPE-Seq (Lucks et al., 2011), MAP-Seq (Seetin et al., 2014), SHAPE-MaP (Siegfried et al., 2014) (Mauger et al., 2015), HRF-Seq (Kielpinski & Vinther, 2014b),Mod-Seq (Talkish et al., 2014), PARS (Kertesz et al., 2010; Wan et al., 2012; Wan et al., 2014), DMS-Seq (Ding et al., 2014; Rouskin et al., 2014), CIRS-Seq (Incarnato et al., 2014), icSHAPE (Spitale et al., 2015). For the studies in which the number of total raw reads were not reported explicitly, plotted values were estimated by *total length * coverage /average read length.* Statistics (blue squares) from EteRNA cloud lab (Lee et al., 2014) involved up to a thousand sequences per round; separate rounds are shown as separate data points. RMDB Accession IDs for datasets shown: (1D). 16SFWJ_STD_0001, TRP4P6_1M7_0006, ETERNA_R80_0001, ETERNA_R82_0001, ETERNA_R83_0003, ETERNA_R86_0000, ETERNA_R87_0003, ETERNA_R92_0000, ETERNA_R93_0000, ETERNA_R94_0000; (2D-M^2^). 16SFWJ_1M7_0001, 5SRRNA_SHP_0002, ADDRSW_SHP_0003, CIDGMP_SHP_0002, CL1LIG_1M7_0001, GLYCFN_SHP_0004, HOXA9D_1M7_0001, RNAPZ5_1M7_0002, RNAPZ6_1M7_0002, RNAPZ7_1M7_0001, RNAPZ12_1M7_0003, TRNAPH_SHP_0002, TRP4P6_SHP_0003; (2D-MaP). adapted from (Homan et al., 2014); (2D-MOHCA). 16SFWJ_MCA_0003, 5SRRNA_MCA_0001, CDIGMP_MCA_0003, GLYCFN_MCA_0002, HCIRES_MCA_0001, HOXA9D_MCA_0001, RNAPZ5_MCA_0001, RNAPZ6_MCA_0002, RNAPZ7_MCA_0001, TRP4P6_MCA_0004; (3D-M^2^-rescue). 16SFWJ_RSQ_0001.

When plotted on a log-log scale and extrapolated to longer RNA lengths (Figure 9B), the strong rise of MCM sequencing costs with number of nucleotides is apparent, especially compared to one-dimensional chemical mapping approaches, which scale linearly with RNA length (blue curve in Figure 9). At the time of writing, four billion reads can be achieved in a large-scale sequencing experiment if all lanes of an Illumina HiSeq machine are put to use (top dashed line, Figure 9B). A single experiment thus allows one-dimensional chemical mapping of most of the highly expressed transcripts in a eukaryotic transcriptome (10^6^-10^7^ nucleotides) (Kwok et al., 2015). For the same cost, MCM methods could, in principle, be applied to a single transcript with a length of at most 10,000 nucleotides. In practice, however this is still overly optimistic. First, available sequencing technologies remain limited to read lengths of a few hundred nucleotides, though this is improving. More fundamentally, all current MCM methods require primer extension by reverse transcriptase to connect events at one nucleotide to a second nucleotide (see, e.g., green arrows in Figures 5A and 5B). With currently tested reverse transcriptases, primer extension is inefficient for lengths beyond 1000 nucleotides even on unmodified transcripts; and, after chemical treatment, modified nucleotides either stop the enzyme (in M^2^ and MOHCA-seq) or reduce its processivity (under conditions tested for RING-MaP/MaP-2D). Based on these considerations, structure determination of a thousand-nucleotide RNA appears to be the upper limit for current MCM methods, and even then would require significant methods development, such as characterization of newly available reverse transcriptases (Mohr et al., 2013). These calculations indicate that application of MCM to infer structures larger than 1000 nucleotides will require a fundamental advance in the methodology. Similar fundamental limitations will prevent application of current MCM methods to model a multitude of transcripts or to uncover intermolecular interactions.

### 6.2 Proposal to overcome sequencing costs

#### Modify-crosslink-and-map

Inspection of current data and new sequencing protocols suggests an experimental strategy to bypass the ∽1000-nucleotide limit to MCM imposed by the quadratic growth of sequencing costs with sequence length. Most of the sequencing reads in an M^2^ or MaP-2D experiment (Figures 3 and 4) correspond to modifications at unstructured nucleotides that are not informative about RNA-RNA contacts (Figure 9A). Even for MOHCA-seq, which focuses its reads on proximal nucleotide pairs, background reverse transcriptase stops at non-proximal nucleotide pairs (which are subtracted from the maps in Figure 5) dominate the sequencing cost for long RNAs (Figure 9). Therefore, any experimental workup that filters out these uninformative hits at unstructured nucleotides and thereby focuses sequencing onto pairs of regions that are roughly proximal could bring the scaling of sequencing costs to be less than quadratic in RNA length. Assuming that each segment of an RNA molecule has a bounded number of possible neighbors within a bounded number of possible states, the sequencing costs would become linear and not quadratic in transcript size.

A method to coarsely filter for proximal segment pairs prior to sequencing can be envisioned by analogy to recently developed crosslinking/sequencing protocols. For example, <u>C</u>ross-linking <u>L</u>igation <u>A</u>nd <u>S</u>equencing of <u>H</u>ybrids (CLASH) and similar approaches (Helwak & Tollervey, 2014) target RNA-RNA interactions by carrying out chemical cross-linking (primarily of nucleic acids bound to proteins), separation of these cross-linked species, removal of unstructured nucleotides through limited nuclease digestion, and ligation of the remaining segments into chimeric sequences (Figure 10B). See also RNA proximity ligation (Ramani et al., 2015), which relies on ligation steps. The ligated segments are then reverse transcribed into chimeric cDNAs for amplification and sequencing; the cross-linked regions are recognized by aligning subsequences against the original transcript or transcriptome sequences (Figure 10C). These methods for inferring nucleotide-nucleotide contacts are powerful for inferring nucleic-acid interactions at the domain level but, in general, their resolutions are too poor for nucleotide-resolution *de novo* structure inference. Ligation boundaries are typically distal to the sites of the crosslinks, and even when mapped through mutational profiling, these nucleotide-nucleotide chemical crosslinks are sparse and can give false positives (Anokhina et al., 2013; Dai et al., 2008; Hang et al., 2015; Levitt, 1969; Robart et al., 2014; Sergiev et al., 2001; Whirl-Carrillo et al., 2002). However, if chemical modifications correlated at a large number of proximal nucleotide pairs are introduced prior to the crosslinking (Figure 10A), they will later give rise to mutations in the final chimeric cDNAs (Figure 10C) upon reverse transcription via the MaP protocol (Homan et al., 2014; Siegfried et al., 2014). The recovery of these correlations induced by single-nucleotide chemical modifications (rather than by crosslinks) would yield rich and accurate MCM measurements, but focused on RNA segment pairs that are roughly proximal *in vivo,* trapped by crosslinking. The cost of this <u>m</u>odify-<u>cross</u>link-<u>m</u>ap method (MXM) protocol would scale in a reasonable manner-linearly with RNA length, if carried out as described above. A series of cross-linking and nuclease digestion times might need to be tested, varied in 2-fold increments to separately capture fragments from easily digested or difficult-to-digest RNA structures. In this case, the scaling of MXM would still increase only loglinearly with RNA length (Figure 9, green dashed line). As a result, MXM should be a viable approach to *de novo* RNA structure characterization for bacterial transcriptomes and for targeted subsets of eukaryotic transcriptomes.

**Figure 10.**
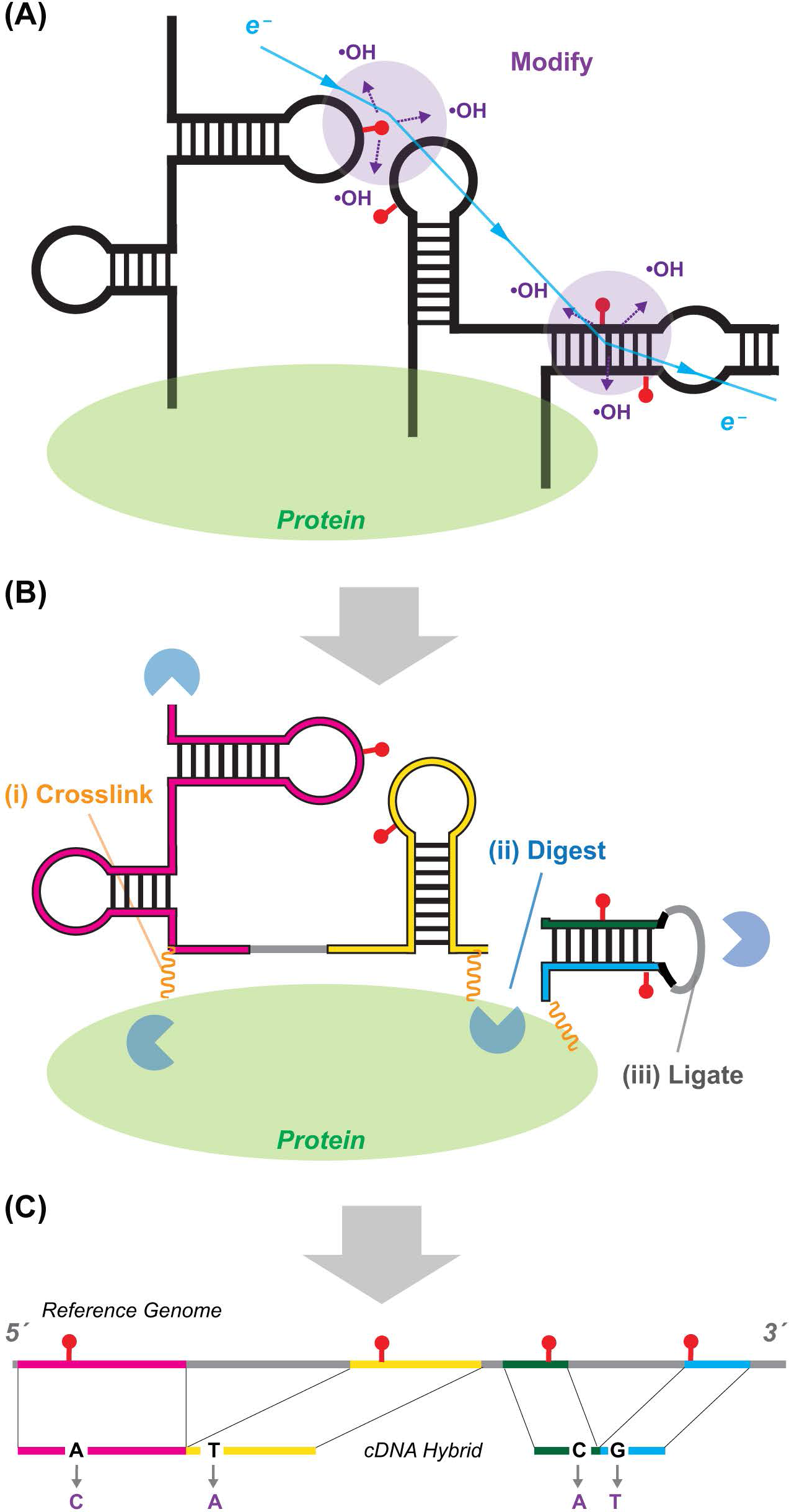
Schematic of the proposed modify-crosslink-map (MXM) expansion. (A). Correlated chemical modifications mark nucleotides brought together by RNA/protein structure *in vivo.* Shown are two sets of oxidative modifications produced by localized ‘spurs’ of hydroxyl radicals generated by scattering of a high-energy electron from water (Chatterjee et al., 1994; Krisch et al., 1991). (B) Additional processing steps of (i) sparse chemical crosslinking, (ii) nuclease digestion, and (iii) RNA ligation (Helwak & Tollervey, 2014) result in compact, chimeric RNA segments harboring correlated chemical modifications. This procedure removes unstructured RNA loops that yield no pairwise structural information and brings together segments distal in sequence or in different RNA strands. (C) Reverse transcription with mutational profiling (Siegfried et al., 2014) reads out modifications at nucleotide resolution; sequence contexts for the modifications allow their alignment to the reference genome sequence.

#### Additional advantages but multiple steps

The MXM protocol would give additional advantages besides reduced sequencing costs. First, by crosslinking and ligating separate RNAs brought together by direct interaction or by colocalization in complexes, MXM would expand MCM to detect pairwise structural interactions across transcripts rather than just within each transcript. Second, MXM would address current inefficiencies in reverse transcription of long RNAs and in sequencing of the associated long cDNAs; the long RNAs would be processed through digestion and ligation into smaller chimeras before sequencing (Figure 10C). Third, MXM could aid in experimentally separating multiple states of RNA transcripts prior to applying the computational deconvolution methods of Section 5. For example, suppose a region of a viral RNA genome forms three distinct local structures-one in the capsid, another while sequestering host factors such as miRNAs, and another while being translated by the ribosome. If these three states give rise to separable crosslinked species or are ligated to different RNA partners (miRNAs, ribosome), MXM maps could determine separate structural maps for the different states.

As with all high-throughput sequencing approaches, turning MXM into a quantitative technique for *de novo* structure inference will require significant investment of time and resources. Optimization and accounting for biases at the many steps – modification, cross-linking, separation, digestion, ligation, reverse transcription, amplification, sequencing, and computational dissection – will each be major challenges. Nevertheless, analogous sequencing approaches of comparable complexity are being developed and numerous groups have recognized steps to bring the methods onto a quantitative footing (Eddy, 2014; Konig et al., 2011; Kwok et al., 2015). Excitingly, rich microscopy data becoming available for actively translating ribosomes (Behrmann et al., 2015) will provide gold standard data to test MXM before its *in vivo* application to RNA messages, long non-coding RNAs, and viral genomes that will be difficult to probe with other methods.

## 7. Conclusion

Multidimensional chemical mapping (MCM) seeks detailed structural information about RNA molecules by measuring how the chemical reactivity of each nucleotide changes in response to perturbations at every other nucleotide. Single-nucleotide perturbations developed in recent years include mutations (mutate-and-map, M^2^), chemical modification (MaP-2D), and radical source attachment (multiplexed ·OH cleavage analysis, MOHCA). MCM experiments provide rich information to guide automated computer modeling methods. In the studies to date, MCM methods have given rich data on secondary and tertiary structure that can be assessed through direct visual inspection. When combined with automated computer modeling, the data have given consistently accurate secondary and tertiary structures at nucleotide resolution. The tests include blind structure prediction trials and modeling of RNA domains for which conventional chemical mapping methods have given incorrect or misleading results. RNA molecules that interconvert between multiple states – which are difficult for or require specialized interrogation with other structural techniques – can be dissected by rapid MCM approaches, albeit with lower resolution than single-structure cases. Incisive validation or falsification can be achieved by high-throughput compensatory rescue experiments for RNA secondary structure, but analogous tests for tertiary structure or ensemble models need to be developed.

The most important frontier for MCM will be the rapid *de novo* structure inference of RNA molecules in their native cellular or viral environments, eventually in a transcriptome-wide manner. Numerous chemical, computational, and sequencing challenges can be foreseen in applying MCM *in vivo.* Nevertheless, there appear to be no fundamental limitations precluding the development of such technologies, particularly if integration with crosslinking can reduce sequencing costs and recover nucleotide-resolution interactions across different RNA strands.

## 8. Acknowledgments

We thank C. Y. Cheng for special assistance in preparing MOHCA-seq figures. We gratefully acknowledge members of the Das laboratory for discussions over several years leading to the perspective outlined herein and the Burroughs Wellcome Foundation (Career Award at the Scientific Interface) and the National Institutes of Health (R01 GM102519) for supporting the writing of the review.

